# Transcriptional Feedback Disruption Yields Escape-Resistant Antivirals

**DOI:** 10.1101/464495

**Authors:** Sonali Chaturvedi, Marie Wolf, Noam Vardi, Kelvin Du, Joshua Glazier, Ruian Ke, Matilda F. Chan, Alan S. Perelson, Leor S. Weinberger

## Abstract

From microbes to cancers, drug-resistant ‘escape’ variants cause significant morbidity and mortality^1–7^. Here we present proof-of-concept that disruption of viral auto-regulatory (feedback) circuits strongly inhibits viral replication and confers an extremely high barrier to the evolution of resistance. Using DNA duplexes, we develop single-molecule ‘feedback-circuit disruptors’ that interfere with transcriptional negative feedback in human herpesviruses (both Herpes Simplex Virus 1 and Cytomegalovirus) thereby increasing viral transcription factors to cytotoxic levels. Feedback disruptors exhibit low-nanomolar to picomolar IC-50’s, reduce viral replication >100-fold in culture and in mice, and synergize with the standard-of-care antivirals. Strikingly, no feedback-disruptor escape mutants evolved over >60 days of culture, in contrast to approved antivirals to which resistance rapidly evolved. Overall, the results demonstrate that molecular targeting of feedback circuitry could yield escape-resistant antivirals, potentially enabling development of a new class of antimicrobials.

## Main Text

The probability of drug-resistant ‘escape’ is driven by various mechanisms including preexisting genetic diversity, the rates of mutation, recombination, and selection, and the effective population size^8,9^. These mechanisms generate substantial intra-host genetic diversity for many viruses, including herpesviruses^10–13^, likely accounting for the significant antiviral resistance observed in clinical settings^4,5^. For example, Herpes simplex virus 1 (HSV-1)—a leading cause of blindness—exhibits resistance to acyclovir (ACV) in ~40% of transplant patients^6^ whereas human herpesvirus 5, cytomegalovirus (CMV)—a leading cause of birth defects and transplant failure—can exhibit resistance to ganciclovir (GCV) in 30–75% of patients^7^. ACV and GCV are nucleoside analogues that terminate viral DNA replication and require phosphorylation by herpesvirus kinases. Resistance arises because single-base kinase mutations ablate kinase activity and such mutations arise relatively rapidly (*μ* ~10^−3^)^11,14,15^ driving the generation and selection of escape mutants^7,16,17^.

Antivirals against alternative targets have been developed^18,19^ but resistance to these new therapies has already been reported^20,21^, consistent with the concept that resistance to antimicrobials is generally unavoidable^1–3^.

Combination therapy, wherein multiple drugs simultaneously inhibit different viral targets, can be employed to limit resistance. For a multi-drug therapy requiring *n* mutations for resistance, escape mutants are predicted to arise at a rate ~*(μ)*^*n*^, requiring substantially larger effective population sizes (i.e., *N* > *(μ)*^−*n*^). However, effective combination therapy requires that each constituent antiviral have a distinct molecular target, and that the combination has favorable toxicity, efficacy, bioavailability, and dosing profiles, which can be challenging^22–24^ (to date, no approved combination regimens are available for herpesviruses).

A proposed alternative to combination therapy has been to mimic its evolutionary benefits by inhibiting protein-protein or protein-DNA interactions^25^ using a single molecule; transcriptional auto-regulatory (feedback) circuits present an attractive target for this approach^26^. Both CMV and HSV-1 utilize transcriptional feedback to regulate immediate-early (IE) viral gene expression, which is obligate to transactivate downstream viral genes, ultimately licensing virus maturation^27–30^. In CMV, the 86-kDa immediate early protein (IE86; a.k.a., IE2), and in HSV-1 the IE175 protein (a.k.a., ICP4), are indispensable transcriptional transactivators^31,32^. Critically, IE proteins can be cytotoxic when expression rises above tightly auto-regulated homeostatic levels, and both CMV and HSV-1 encode negative-feedback circuits to maintain IE86 and IE175 levels below their respective cytotoxic thresholds^31,33–35^. These feedback circuits are comprised of a protein-DNA interaction wherein the IE protein binds to a 14–15 bp palindromic cis-repression sequence (crs) within its respective promoter and auto-represses its transcription (Fig. 1a). Previous studies showed that disrupting IE feedback, via mutation of the crs, increases IE protein levels past a cytotoxic threshold and leads to a >100-fold reduction in viral replication^33,36,37^. Building off the concept of transcription factor decoys^38^, we hypothesized that short double-stranded DNA oligonucleotide duplexes that mimic the crs DNA-binding site could act as competitive inhibitors to disrupt IE negative feedback, thereby increasing IE levels into the cytotoxic regime (Fig. 1a). Mathematical modeling predicted that such feedback disruptor (FD) molecules could in principle raise IE protein to cytotoxic levels (Fig. 1b and Extended Data Fig. 1a). To estimate the likelihood of resistance to FDs arising, we used computational models previously used to calculate GCV resistance^7^ and calculated the probability that pre-existing mutants could undergo selection (Fig. 1c). Theoretically, resistance to an FD would require the virus to recapitulate a new negative-feedback loop, which would require evolution of an orthogonal DNA-binding domain in the protein (to recognize a new orthogonal DNA sequence) and *simultaneous* evolution of the new orthogonal DNA-binding sequence. A substantial body of literature has shown that engineering of altered DNA-protein binding specificity requires 30–50 mutations distributed between the DNA and protein^25,39,40^ and our calculations (Fig. 1c) indicate that the frequency of pre-existing mutants scales as ~*μ*^−*n*^ such that FD-escape mutants (*n* > 30) have a vanishingly small probability, whereas ACV/GCV resistance mutants, which require only a single mutation, arise with high probability. While it is conceivable that mutants in the IE protein could arise and bind alternate crs-like sequences in the promoter region, such mutants were not observed in month-long culturing^33^.

**Figure 1:**
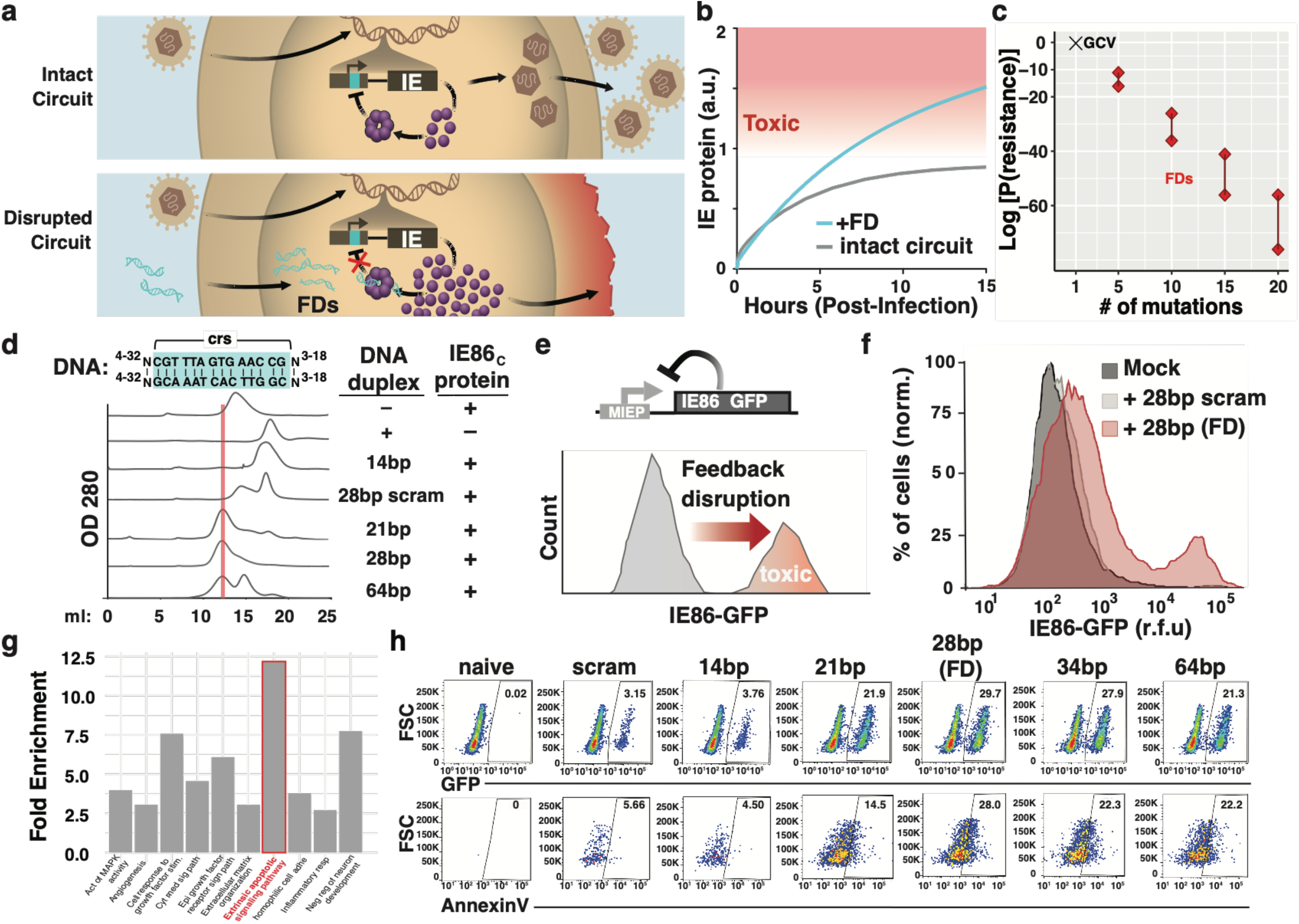
Identification of feedback disruptor DNA duplexes. (a) Schematics of the herpesvirus IE (IE86 and IE175) transcriptional negative-feedback circuits in the intact wild-type form (upper) and after disruption (lower) by putative feedback disruptors (FDs). When feedback is intact, IE proteins bind the cis repression sequence (crs) in their respective IE promoters (cyan) and downregulate transcriptional activity to prevent IE protein levels from reaching cytotoxic levels. When feedback is disrupted, for example by IE proteins being titrated away by binding free double-stranded (ds) DNA duplexes encoding the crs, IE promoter activity is not downregulated and IE proteins reach cytotoxic levels (~1.5-fold above homeostatic levels)^33^. (b) Numerical solutions of an experimentally-validated computational model of IE feedback^33,43^ showing that FDs effectively break feedback to increase IE protein levels to the cytotoxic regime. See also Extended Data Fig. 1a. (c) Probability of a drug-resistant variant arising during the course of therapy (y-axis). For GCV (‘x’), 1–2 mutations are needed to generate resistance and the probability of generating resistance is between 70%–100%. However, for FD resistance, multiple mutations are needed and the probability of a resistant variant arising becomes negligibly small. The line between the diamonds shows the uncertainty ranges arising from the uncertainty in the mutation rate estimated as 10^−3^–10^−4^). See methods for details of the calculation. (d) Top: Schematic of different lengths of DNA duplexes tested for oligomerization of the IE86 protein (c-terminus fragment). Bottom: Chromatographs of c-terminus of IE86 fragment incubated with either a sequence-scrambled control DNA duplex or crs-containing DNA duplexes of differential lengths. The 28bp crs DNA duplex most efficiently titrates free protein from the 15mL fraction into the 13mL fraction (~98% of protein is found in the 13mL protein-DNA complex fraction when the 28bp crs DNA duplex is added). (e) Schematic of the minimal IE negative-feedback circuit (MIEP-IE86-IRES-GFP) that is contained within the feedback-reporter cell line and schematic of how negative-feedback disruption increases GFP fluorescence in the feedback-reporter cell line. (f) Flow cytometry of feedback-reporter cells 48 h after nucleofection with either 28bp DNA duplex encoding a crs, a scrambled DNA duplex (negative control), or mock nucleofection (no DNA duplex) showing that the crs-encoding 28bp DNA duplex disrupts feedback. (g) RNAseq analysis of feedback-reporter cells at 72 h post feedback disruption with 28bp DNA compared to nucleofection of scramble 28bp scramble. Gene ontology (GO) analysis of enriched pathways plotted for the top ten most-enriched pathways. (h) Flow cytometry of feedback-reporter cells 72 h after nucleofection with DNA duplexes (see Table 1 for further information) comparing circuit disruption vs. early apoptosis, by IE86 expression (GFP) and Annexin V, respectively (see Extended Data Fig. ** for TUNEL staining). Values within the plots report % positive cells in the respective gate. The 28bp DNA duplex exhibits maximal feedback disruption (% YFP+ cells) and apoptosis induction (% Annexin V+ cells), and is designated a feedback disruptor (FD).

To find double-stranded DNA oligonucleotides (duplexes) that optimally titrate IE proteins, we developed an *in vitro* liquid-chromatography assay to quantify the efficiency of various linear DNA duplexes in catalyzing formation of the IE86 protein-DNA complex (Fig. 1d). To validate the assay, we used electrophoretic mobility shift assays (EMSA) and verified that purified IE86 protein bound to DNA duplexes in a sequence-specific manner (Extended Data Fig. 1b). We tested an array of crs-encoding duplexes of various lengths and found a linear 28 base-pair (bp) DNA duplex most efficiently catalyzed formation of the IE86-DNA complex. Shorter or longer crs-encoding DNA duplexes were less efficient at promoting protein-DNA complex formation (Fig. 1d, Extended Data Fig. 1c). We confirmed that the protein-DNA oligomerization was not a result of the maltose binding protein (MBP) tag at the n-terminus of the purified protein (Extended Data Fig. 1d). Since the purified IE86 protein consisted of the C-terminus of IE86, we verified that the 28bp-DNA duplex also catalyzed protein-DNA oligomerization for the full-length IE86 protein (Extended Data Fig. 2a).

To test if these DNA duplexes disrupted transcriptional negative feedback, we used a cell line stably transduced with a minimal IE86 negative-feedback reporter circuit^33^. In these retinal pigment epithelial cells, IE86 and GFP are expressed from the IE86 promoter-enhancer and are subject to IE86 auto-repression. Increases in GFP indicate disruption of negative feedback and lead to cell death within 48–72 hours after feedback is disrupted (Fig. 1e). These reporter cells were nucleofected with DNA duplexes—stabilized against nuclease degradation by internal phosphorothioate bonds^41,42^ (Extended Data Fig. 1f)—and cellular uptake of DNA duplexes was verified by Cy3 fluorescence (Extended Data Fig. 1e). The 28bp-DNA duplex substantially disrupted IE86 feedback, increasing GFP expression and cell death, whereas the 28bp scrambled control did not disrupt feedback or increase cell death (Fig. 1f). Feedback-circuit disruption was dose-dependent (Extended Data Fig. 1g, h) and was enhanced by duplex concatemerization, whereas a non-duplex, single-stranded DNA oligonucleotide control, with the same 28-base sequence, did not disrupt feedback (Extended Data Fig. 2e, f, g).

To determine the mechanism by which feedback disruption induced cell death, we performed RNAseq analysis (Fig. 1g) to identify upregulated cell-death pathways. Differential transcriptome analysis indicated a number of overexpressed pathways in feedback-disrupted cells, relative to cells nucleofected with the scrambled 28bp DNA control, with apoptotic pathways prominently enriched (Fig. 1g). Apoptotic pathway enrichment was confirmed by a second gene-ontology algorithm (Extended Data Fig. 3). To verify that apoptosis was upregulated in cells with disrupted IE86 feedback, we stained for both the early and late apoptotic markers (Fig. 1h, Extended Data Fig. 1i). In agreement with the size-exclusion chromatography above, feedback disruption was dependent on DNA-duplex length: while the 34bp and 64bp-DNA duplexes showed some efficacy, the 28bp-DNA duplex (FD^C^) exhibited maximum feedback disruption (i.e., max increase in %IE86-GFP+ cells) and maximum Annexin V staining (Fig. 1h). These data were confirmed via a second assay (Extended Data Fig. 1i), Terminal deoxynucleotidyl transferase (TdT) dUTP Nick-End Labeling (TUNEL), which detects DNA degradation during the late stage of apoptosis. Collectively, these results indicate that relatively short (< 30bp) sequence-specific DNA duplexes are capable of viral feedback-circuit disruption leading to cytotoxicity and we thus designated the 28-bp DNA duplex an IE86 feedback disruptor for CMV (FD^C^).

We next investigated the effects of disrupting IE86 feedback in the context of viral infection. We first confirmed that the FD^C^ DNA duplex did not interfere with viral entry, using two separate assays (Extended Data Fig. 4a–c). Flow cytometry and microscopy analysis showed that the FD^C^ DNA duplex disrupted IE86 negative feedback and increased IE86 expression >10-fold in cells infected with a CMV TB40E virus carrying a IE86-YFP fusion^43^ (Fig. 2a). Dose-response analysis of FD^C^‘s effect on single-round viral replication showed an IC-50 = 0.95 nM (Fig. 2b), with ~100-fold repression of viral replication (in agreement with genetic disruption of IE negative feedback^33^). This inhibition of viral replication was consistent across a broad range of multiplicity of infection (MOI) from 0.1–2.0 (Fig. 2c). To be sure that the observed antiviral effects were not specific to the virus strain or cell type used, we also tested (i) CMV strain AD169 and (ii) various GCV-resistant CMV strains in human foreskin fibroblasts, as well as (iii) murine CMV and (iv) rhesus CMV, in mouse and primate cells respectively, and in all cases found similar 100-fold viral titer reduction using FDs designed for the respective viral crs sequences (Extended Data Fig. 5a–d).

**Figure 2:**
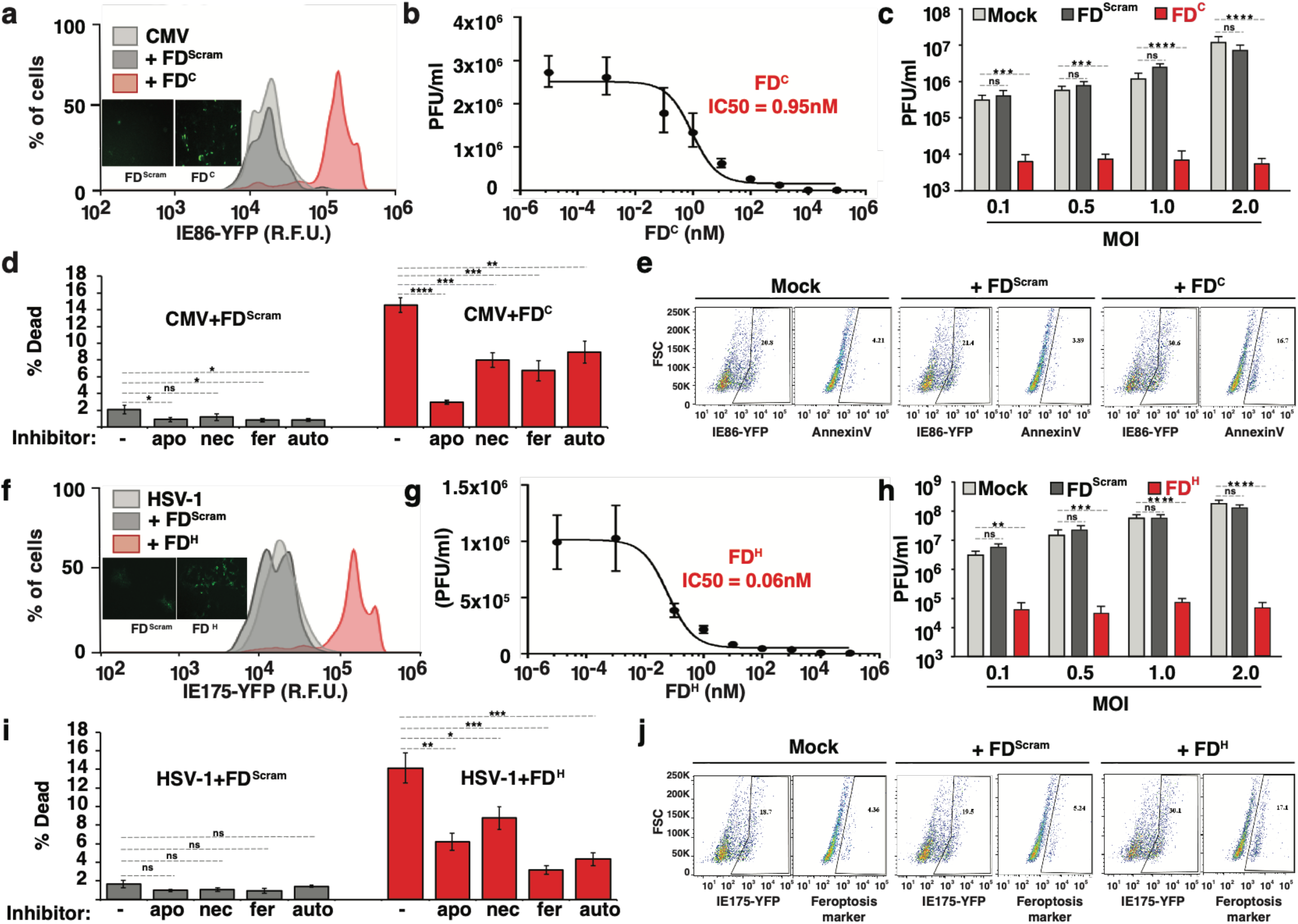
Feedback disruption interferes with viral replication by inducing cytotoxicity in infected cells. (a) Flow cytometry analysis showing that feedback disruption generates IE86 overexpression in CMV-infected cells. Retinal pigment epithelial (ARPE-19) cells were nucleofected with the 28bp DNA duplex that disrupts CMV (FD^C^) at 25μM or a scrambled DNA duplex (FD^Scram^), infected with a CMV (TB40E) encoding an IE86-YFP fusion (MOI = 0.1) and analyzed at 2 days post infection (dpi). Insets: Micrographs of YFP fluorescence in TB40E IE86-YFP infected ARPE-19 cells at 24 h post infection (MOI=1.0). (b) FDC dose-response curve and corresponding IC_50_ value. ARPE-19 cells were nucleofected with FD^C^ at concentration specified, infected with TB40/E-IE86-YFP virus (MOI=0.1), and virus titered 4 days later. (c) Viral knockdown by FD^C^. ARPE-19 cells were nucleofected with 25μM FD^C^ (or mock/FD^Scram^), and 24 hours post nucleofection, cells were infected with TB40/E-IE86-YFP virus at different MOIs (0.1, 0.5, 1.0, 2.0), and titered at 4 dpi. (d) Cytotoxocity ‘rescue’ assay showing FD^C^ induces apoptosis CMV-infected cells. ARPE-19 cells were nucleofected with 25μM FD^C^ (or mock/FD^Scram^) and infected with TB40/E-IE86-YFP virus at 24 h post nucleofection in the presence of the indicated cell death inhibitors: *auto* (authophagy inhibitor); *apo* (apoptosis inhibitor); *nec* (necroptosis inhibitor); *fer* (ferroptosis inhibitor). Cells were harvested at 48 hpi and stained for dead cells with Zombie Aqua (BioLegend). (Experiment performed in three biological replicates; p values calculated using 2-way ANOVA followed by Tukey’s multiple comparisons test). (e) Apoptosis induction in CMV-infected cells: ARPE-19 cells were nucleofected with 25μM FD^C^ (or mock/FD^Scram^), 24 hours later cells were infected with CMV (TB40/E) IE86-YFP virus (MOI=1), and at 48 h post infection stained with Annexin V. (f) Feedback disruption leads to IE175 overexpression in HSV-1 infected cells. Cells (ARPE) were nucleofected with the 29bp DNA duplex that disrupts HSV-1 feedback (FD^H^) at 25μM FD^H^ (or mock/FD^Scram^), infected with HSV-1 (17syn+ strain) encoding an IE175-YFP (MOI = 0.1) and analyzed at 2 dpi. Insets: Micrographs of Vero cells infected with 17syn+ IE175-YFP at 12 h post infection (MOI=1.0). (g) FDH dose-response curve and corresponding IC_50_ values. Titers were calculated on Vero cells nucleofected with FD^H^ at concentration indicated, infected with HSV-1 IE175-YFP (17syn+ strain, MOI=0.1) 24 h later and tittered 2 days post infection. (h) Vero cells were nucleofected with 25μM FD^H^ (or mock/FD^Scram^), and 24 hour post nucleofection, cells were infected with HSV-1 (17syn+ strain) at different MOIs (0.1, 0.5, 1.0, 2.0), and titered at 4 dpi. (i) Cytotoxocity ‘rescue’ assay: FD^H^ induces ferroptosis in HSV-infected cells. Vero cells were infected with HSV-1 (17syn+ strain) IE175-YFP virus (MOI 0.1) at 24 hour post nucleofection with FD^Scram^ or FD^H^ in the presence of the indicated cell death inhibitors. Cells were harvested at 48 hpi and stained for dead cells with Zombie Aqua (BioLegend), and subjected to flow-cytometry. Experiment performed in three biological replicates. p values calculated using 2-way ANOVA followed by Tukey’s multiple comparisons test. (j) Ferroptosis induction in HSV-infected cells Vero cells. Vero were nucleofected with 25μM FD^H^ (or mock/FD^Scram^), at 24 hour post nucleofection, cells were infected with HSV-1 (17syn+ strain) IE175-YFP (MOI=1), and at 24 hour post infection cells were harvested and stained with a ferroptosis marker. p-values less than 0.05 were considered statistically significant: *<0.05, **<0.01, ***<0.001, ****<0.0005, ns= not significant.

To determine whether IE86 feedback disruption and overexpression also led to apoptosis in the context of viral infection, we used a cell-death rescue assay that employs inhibitors of various cell-death pathways (Fig. 2d, Extended Data Fig. 6)—RNAseq of infected cells was dominated by response-to-infection pathways and was unable to discern cell-death pathways (not shown). The rescue assay indicated that IE86 feedback disruption during CMV infection activates many cell-death pathways—possibly due to crosstalk between cell-death pathways^44^—but apoptosis inhibition gave the most pronounced minimization of cell death in FD-treated CMV-infected cells (Fig. 2d). Staining for apoptosis in CMV-infected cells confirmed that treatment with FD^C^ increased apoptosis, whereas treatment with the sequence-scrambled duplexes (FD^Scram^) did not increase apoptosis in infected cells (Fig. 2e).

Based on feedback disruption inhibiting viral replication in diverse CMVs (β herpesviruses), and findings that HSV-1 (an α-herpesvirus) encodes a similar negative-feedback circuit^37^, we hypothesized that replacing the 14bp crs within FD^C^ with the 15bp repression sequence from the HSV-1 IE175 promoter could generate a FD for HSV-1 (this 29bp-DNA duplex was designated FD^H^). As predicted, FD^H^ disrupted IE175 negative feedback in cells infected with the 17-syn+ HSV-1 strain^45^ and increased IE175 expression >10-fold (Fig. 2f), without altering viral entry (Extended Data Fig. 4 b, c). Strikingly, FD^H^ exhibited an IC-50 = 0.06 nM (Fig. 2g) and inhibited HSV-1 replication ~100-fold across a broad range of MOIs from 0.1–.0 (Fig. 2h). Analysis of cell-death pathways in the context of HSV-1 infection showed that ferroptosis inhibition minimized FD^H^-induced cell death, though other cell-death pathways also appear to play a role (Fig. 2i, j, Extended data Fig. 6). Similar antiviral effects were observed for a second HSV-1 strain (KOS) (Extended data Fig. 5e), and these results were observed in both Vero and ARPE-19 cells (Fig. 2h, 3h).

Importantly, sequence-scrambled duplexes (FD^Scram^) did not exhibit antiviral effects, suggesting that FDs do not act nonspecifically via innate-immune mechanisms (e.g., activation of cGAS-STING pathway via TLR9), which is consistent with efficient cGAS pathway activation requiring DNA duplexes of > 300bp^46^ (whereas FDs are < 30bp). However, to verify that FD activity is independent of cGAS-STING, we tested FDs under conditions of high and low cGAS-STING expression and observed little difference in FD antiviral effects and no effects for FD^Scram^ in either setting (Extended Data Fig. 7e). Moreover, FDs did not activate TLR9 expression (Extended Data Fig. 7d).

To confirm the mechanism of action—that FD-induced cell death was due to disruption of viral negative feedback—we used a previously-characterized mutant virus^33^ which lacks IE86 negative feedback due to a two-base mutation in the crs region of the promoter (Fig. 3a). As expected, FD^C^ did not reduce viral replication of the mutant virus (Fig. 3b) or significantly increase IE86 expression in this mutant (Fig. 3c). These data confirm that inhibition of CMV viral replication by FD DNA duplexes is due to specific disruption of IE86 transcriptional feedback circuit.

**Figure 3:**
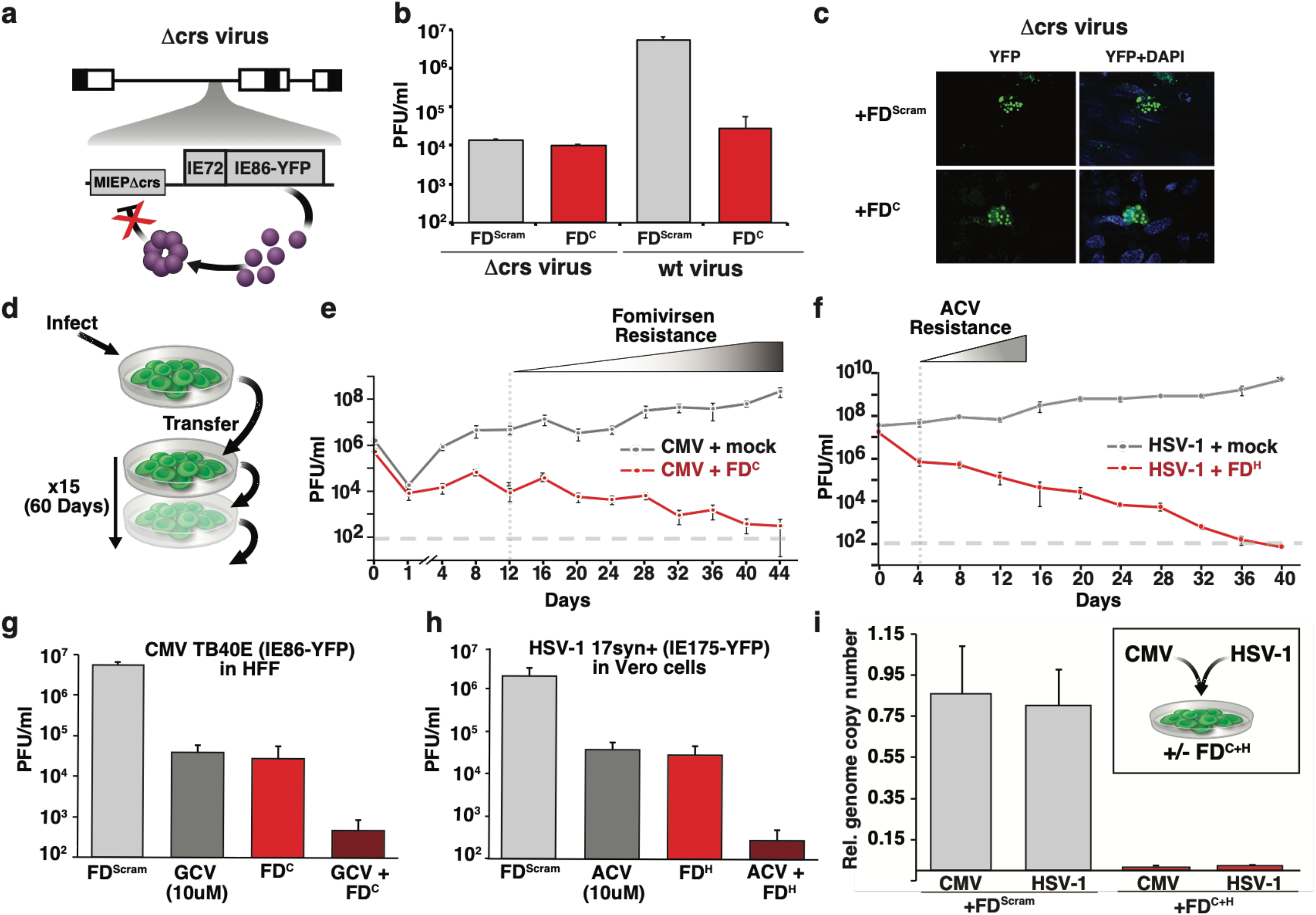
Feedback disruptor mechanism, recalcitrance to viral escape, and combinatorial activity. (a) Schematic of the Δcrs CMV virus that has disrupted IE86-feedback feedback due to a 2bp-mutation (CG**→**AA) in the crs region of the promoter where IE86 binds. (b) FD^C^ does not reduce Δcrs virus replication. Cells (human foreskin fibroblast, HFF) were nucleofected with 25μM FD^C^ (or FD^Scram^) and 24-hour post nucleofection infected with parent AD169 CMV virus or Δcrs AD169 CMV mutant virus (MOI=0.1) and then titered (4-dpi for parent virus and 6-dpi for Δcrs virus due to its slower growth). (c) FD^C^ does not increase IE86 levels Δcrs virus infected cells. Fluorescent micrographs of HFF cells infected with Δcrs virus in the presence of FD^Scram^ or FD^C^ (6 days post infection). (d) Schematic of the continuous-culture experiment; ARPE-19 cells (+/− FD) were infected with CMV or HSV-1 (MOI=0.1) and at 4-dpi, supernatant was collected and was used to infect naïve ARPE-19 cells (+/− FD) until day 60. (e) Continuous culture titers for CMV (TB40E-IE86-YFP) in the presence of FD^C^ (red) or mock treatment (grey). Fomivirsen resistance (positive slope of the titering dynamics) was observed beginning at day 12 (Extended Data Fig. 7f-h). (f) Continuous culture for HSV-1 (17syn+ IE175-YFP virus) in the presence of 25μM FD^H^ (red) or mock treatment (grey). ACV resistance (positive slope of the titering dynamics) observed beginning at day 4 (Extended Data Fig. 7i). (g–h) Combinatorial effect of FDs with standard-of-care antivirals for CMV and HSV-1. (g) Virus titer for CMV in the presence of FD^Scram^, ganciclovir (GCV, 10μM), 25μM FD^C^ or (10μM GCV and 25μM FD^C^) in HFF cells at 4 dpi. (h) Virus titer for HSV-1 in the presence of FD^Scram^, acycolovir (ACV, 10μM), 25μM FD^H^ or (10μM ACV and 25μM FD^H^) in Vero cells at 2 dpi. (i) Multiplexed feedback disruption can inhibit CMV and HSV-1 replication. Inset: Schematic of the mixed-infection experiment; ARPE-19 cells nucleofected with equimolar amounts of FD^C^ and FD^H^ or FD^Scram^ were co-infected with CMV (TB40E-IE86-YFP) and HSV-1 (17syn+ IE175-YFP) at MOI = 0.1. Main: qPCR analysis of CMV and HSV-1 viral genomes in a mixed-infection experiment at 4 dpi using primers specific for CMV and HSV-1 (Bar graph) (see Extended Data Table 1 for sequences).

To test the hypothesis that feedback disruption presents a high barrier for the evolution of resistance, we employed both phenotypic and genotypic resistance assays. For phenotypic resistance assay, we developed a continuous-culture approach (Fig. 3d) wherein virus was consecutively passaged from infected cells to fresh uninfected cells every 4 days (a typical CMV replication round, >4 HSV-1 replication cycles). These continuous cultures were carried until virus was undetectable in the presence of FD treatment (~40–60 days; Fig. 3e, f). The FD^C^ antiviral effect was benchmarked against Fomivirsen^47^, the first approved oligonucleotide therapy (anti-sense DNA for IE86) (Fig. 3e), whereas FD^H^ was compared to ACV (Fig. 3f). As previously reported^16,17^, we found HSV-1 resistance to ACV emerged within two rounds of infection (Extended Data Fig. 7i) and that CMV resistance to Fomivirsen^47^ emerged within 3-4 rounds of infection (Extended Data Fig. 7g, h). In stark contrast to these approved antivirals, FD^C^ steadily reduced CMV titers to below the limit of detection by day 52, with no evidence of CMV resistance to the FD^C^ (Fig. 3e). Subsequent sub-culturing in the absence of FD^C^ showed that the virus was cleared (Extended Data Fig. 7f). Similarly, FD^H^ steadily reduced HSV-1 titers to below detection by day 40, with no evidence of resistance (Fig. 3f). For the genotypic resistance assay, sequence analysis could not detect mutations anywhere in the 500bp-region surrounding the promoter repression sequences (Extended Data Fig. 8), and only detected transient single-nucleotide polymorphisms in the IE-protein regions responsible for DNA binding (Extended Data Fig. 9). Overall, these results support the hypothesis that feedback disruption has a high barrier to the evolution of resistance.

When feedback disruptors were tested together with the standard-of-care therapies, GCV (Fig. 3g) or ACV (Fig. 3h), FDs generated reductions in viral titer comparable to GCV and ACV, but, strikingly, combinations of FD^C^+GCV and FD^H^+ACV generated additive reductions in viral titer of ~4-Logs, consistent with FDs acting via a mechanism distinct from GCV and ACV. To test if FDs could be multiplexed—given that herpesvirus infections of unknown etiology are a significant clinical problem^48,49^—cells were co-transfected with FD^C^ and FD^H^, or FD^Scram^ alone as a control, and then infected with both CMV and HSV-1 (MOI 0.1). qPCR analysis showed that relative genomic copy number of both CMV and HSV-1 was effectively reduced (Fig. 3i). The results suggest that FDs could be used in conjunction with standard-of-care therapies to enhance the antiviral effect and further minimize resistance, and could be multiplexed.

Finally, we tested if transcriptional feedback could be disrupted *in vivo* using the established model of herpes infection in mice^50^. In this model, mice are infected with HSV-1 in the cornea, and interventions are topically applied at the site of infection. We infected mice with HSV-1 IE175-YFP, then applied duplexes 6 hours later. After two days, corneas were harvested for imaging and quantification of viral replication by q-PCR and titering (Fig. 4a). FD^H^ uptake by cells was observed by Cy3 fluorescence (Extended Data Fig. 5f) and as predicted, FD^H^ treatment caused an increase in IE175 (Extended Data Fig. 5g) and a significant reduction in the percentage of HSV-1 infected cells (Fig. 4b-c). In agreement with these data, FD^H^ treatment reduced viral titer by 150 fold (Fig. 4d) and significantly reduced viral genome replication (Fig. 4e). Together, these results demonstrate that feedback disruption reduces viral replication *in vivo*.

**Figure 4:**
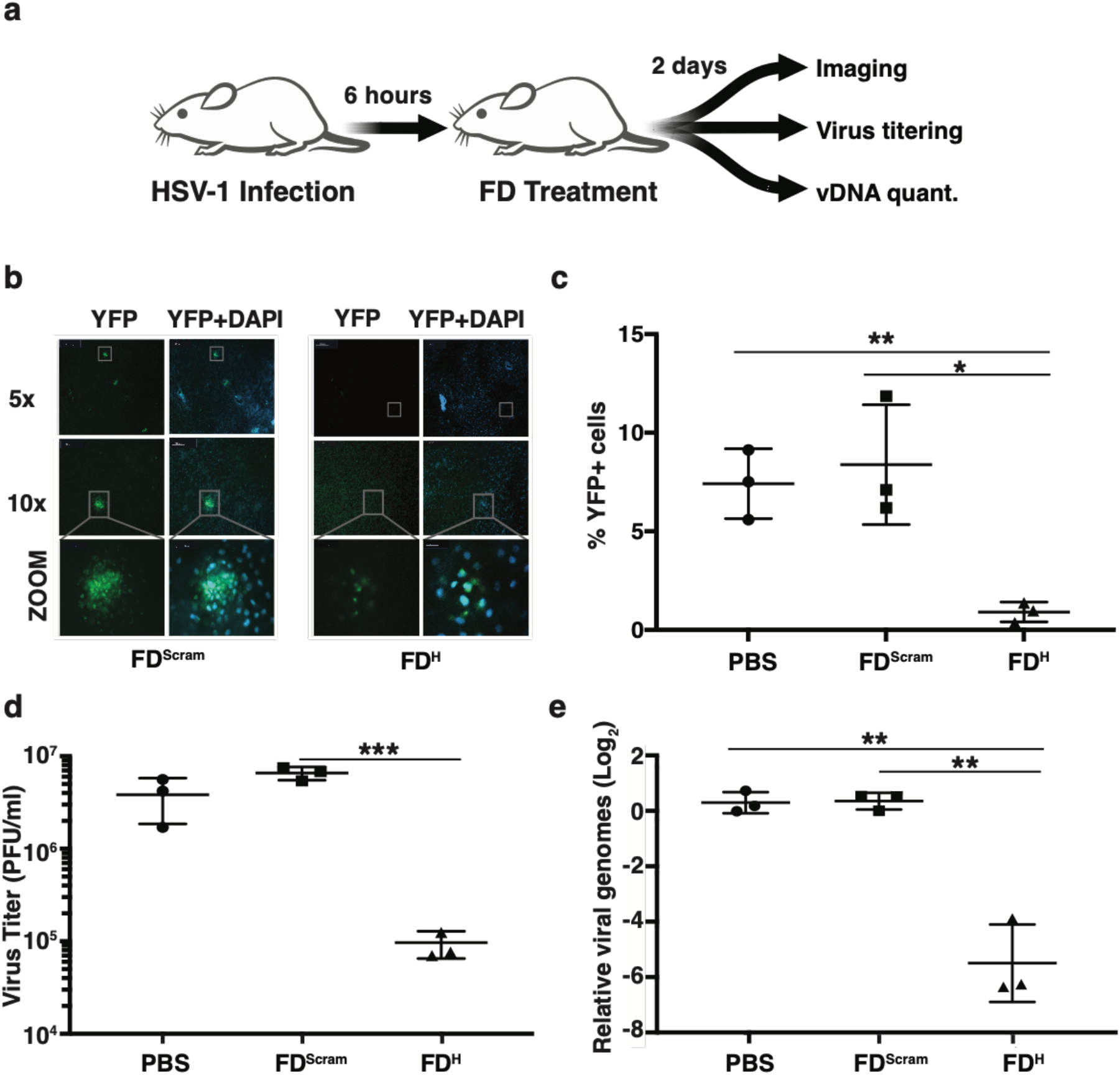
Feedback disruption inhibits viral replication in an *in vivo* model. (a) Schematic of the HSV-1 corneal infection model in mice. BL-6 mice, 6–10 weeks old, undergo corneal debridement followed by infection with HSV-1 17syn+ IE175-YFP virus (1×10^5^ PFU). 6 hours post infection, 25μM FD^H^, or FD^Scram^, or PBS, was topically applied to the cornea. Corneas were harvested at 2 dpi, imaged for YFP, HSV-1 levels quantified by virus titering, and viral genomes quantified by qPCR. (b) Representative YFP-fluorescence images of corneas after harvesting (nuclei stained with DAPI). (c) Quantification of HSV-1 YFP expressing cells in corneas, as determined by masking on DAPI and scoring YFP-positive cells. 5 corneas imaged per sample. p-values derived from Students T test: *<0.05, **<0.01, ***<0.001. (d) HSV-1 viral titers from infected corneas 2 days after treatment with either 25μM PBS, FD^Scram^ or FD^H^. Corneas were dissociated using collagenase, subjected to three freeze-thaw and supernatant was used to titer virus using end time dilution method (TCID50). Each data point represents a pooling of corneas (i.e., 9 corneas per treatment) of two independent experiments. p-values derived from Students T test: *<0.05, **<0.01, ***<0.001. (e) HSV-1 viral genomic DNA quantification by qPCR 2 days after treatment. Each data point represents a pooling of corneas (i.e., 9 corneas per treatment) of two independent experiments. p-values derived from Students T test: *<0.05, **<0.01, ***<0.001.

While delivery of oligonucleotides remains a major challenge, significant clinical advances have been made with the recent FDA approval of antisense and exon-skipping oligonucleotide therapies delivered via nanoparticles^51–53^. Therefore, we also tested FDs using conventional gold nanoparticle carriers^54^, in the absence of transfection, and found similar antiviral effects (Extended Data Fig. 7j, k).

Overall, these results present proof-of-principle that transcriptional feedback could represent a novel antiviral target with a high barrier to the evolution of resistance (Fig. 1c and 3e, f). The results herein also indicate that feedback disruptors could overcome a current treatment challenge: the reduction in antiviral efficacy at high-viremic loads^55^. Robust antiviral effects are notoriously difficult to achieve under high-viremic conditions which increase the potential for outgrowth of resistance mutations. For FDs, however, higher MOIs result in more potentially cytotoxic IE protein in the cell. From the resistance perspective, not only are multiple mutations required to recapitulate the cis-trans protein-DNA interactions required for feedback^25,40^ (Fig. 1c) but viral regulatory loci appear to have lower genetic variability than viral enzymes or receptors^10,12,14^. As such, we predict that feedback circuit disruption may be an attractive target in other viruses, microbes, and possibly neoplasms with aberrant auto-regulatory circuits.

## Abbreviations

(FD): feedback disruptor
(CMV): human cytomegalovirus
(HSV-1): herpes simplex virus 1

## Acknowledgments

We dedicate the manuscript to the memory of Cynthia Bolovan-Fritts, who provided extensive technical support and made this work possible. We thank Lewis Lanier, Peter Barry and Roger Everett for providing MCMV strain K181, Rhesus CMV 68.1 EGFP, HSV-1 17syn+ IE175-YFP virus. We thank Mike Pablo for his mathematical modeling expertise, Elena Ingerman, Melanie Ott, JJ Miranda, Siyu Zheng, Marielle Cavrois, Nandhini Raman, Elizabeth Tanner and the Weinberger lab for discussions and suggestions. We thank Kathryn Claiborn for reviewing the manuscript. We acknowledge Gladstone Flow Cytometry Core, funded through NIH P30 AI027763. We acknowledge the Gladstone Genomics Core and Bioinformatics Core for technical expertise on RNAseq. We acknowledge the James B. Pendelton Charitable Trust for the FACSAria cell sorter. M.F.C. acknowledges National Institutes of Health (R01 EY022739, NIH-NEI EY002162 - Core Grant for Vision Research), Research to Prevent Blindness (RPB Physician-Scientist Award to M.F.C and RPB Unrestricted Grant to the UCSF Department of Ophthalmology), and That Man May See Research Grant. Portions of this work were done under the auspices of the US Department of Energy under contract 89233218CNA000001 and supported by NIH grants R01-OD001095 and R01-AI028433 (to A.S.P.) This work was supported by the Bowes Distinguished Professorship, the Alfred P. Sloan Research Fellowship, the Pew Scholars in the Biomedical Sciences Program, NIH Director’s New Innovator Award OD006677 and Pioneer Award OD17181 programs (to L.S.W.).

## Author contributions

S.C. and L.S.W. conceived and designed the study. S.C., M.W., and M.F.C., designed and performed the experiments, and curated the data. N.V., L.S.W., R.K., and A.S.P performed the mathematical modeling. S.C., M.W. and L.S.W. wrote the paper.

## Author information

Reprints and permissions information is available at www.nature.com/reprints

## MATERIAL AND METHODS

### Feedback Disruptor Mathematical Modeling and Numerical Simulations

We used an experimentally validated ODE model of the CMV IE86 negative feedback circuit^22, 24^ modified to include state variables for free IE86 protein (IE86), crs feedback-disruptor (*FD*) DNA duplexes, and the IE86–FD DNA-protein interaction complex (*Complex)*:

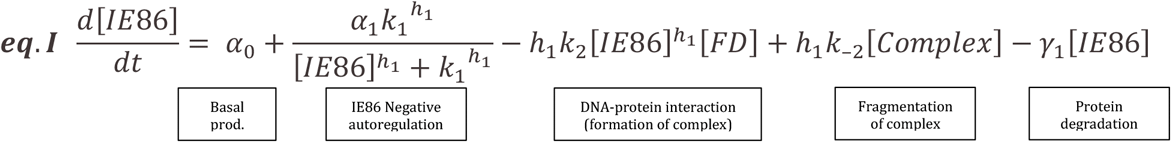

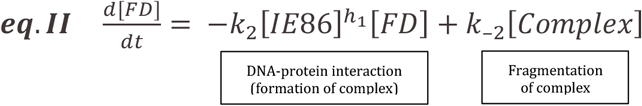

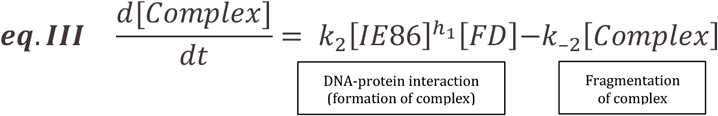

where *α*_0_ represents the basal IE86 expression rate (2 hr^−1^), *α*_1_ represents the IE86 negative-feedback gain constant (10 hr^−1^), *h*_1_ represents the IE86 cooperativity index (hill coefficient), *k*_1_ represents a Michaelis constant (set to 1) for IE86 feedback, *γ*_1_ represents the per-capita IE86 protein degradation rate (0.23 hr^−1^), *k*_2_ represents the *Complex* association/formation rate (2 hr^−1^), and *k*_−2_ represents the *Complex* dissociation/fragmentation rate (1/hr^−1^). In this model, the IE86 can oligomerize (*h*_1_ ~ 6)^22^ to bind either the crs in its own promoter (MIEP)—thereby mediating negative feedback and down-regulating its own expression rate—or can bind the FD DNA duplex, thereby sequestering IE86 that might otherwise downregulate its own expression via negative feedback. For both binding events, IE86 can transition between free and FD-bound states but the negative feedback bound state is not explicitly modeled for parsimony and to not introduce unnecessary parameters. The DNA duplexes were assumed to be stable (i.e., not degrade) over the course of the simulation time (validated below). For Extended Data Fig. 1, the initial levels of FD at time=0 were varied as indicated, all other initial conditions were zero. The ODEs were numerically solved using Matlab™.

### Within-host CMV model for calculation of probability of resistance to FDs

To calculate the probability of mutants arising during infection, we construct a within-host model for CMV infection, based on a previous model widely used for chronic viral infections such as HIV and HCV^8^. The ODE system describing within-host infection dynamics are:

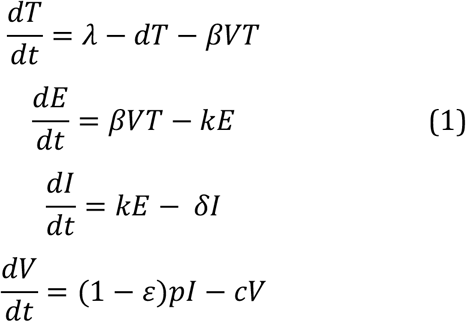

This model considers the dynamics of target cells (*T*), cells infected but not yet producing virus, i.e. in the eclipse phase (*E*), productively infected cells (*I*) and virus (*V*). Target cells are generated at rate, *λ*, and die at per capita rate *d*. They can be infected by virus interacting with target cells at rate *β*, leading to productively infected cells (*I*). Productively infected cells die at a per capita rate *δ* and produce viruses at rate *p* per cell. Viruses are cleared at per capita rate *c*. *ε* models the efficacy of a therapy. Here we assume that a therapy reduces viral production from infected cells. Note that the calculation of probability of resistant mutant appearing in the virus population is robust against this assumption, i.e. if a drug increases the death rate of infected cells, as for FDs, the conclusion remains the same. From Eq. 1, one can calculate the within-host reproductive number, *R*_0_, using the next-generation matrix method ^56^, in the absent of treatment:

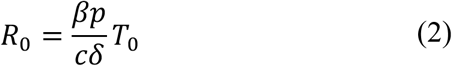

where *T*_0_ is the target cell concentration in an uninfected individual.

Parameter values were chosen similar to other mathematical modeling studies^57^ that examine the action of ganciclovir (GCV) on intracellular CMV replication, which estimate the infected cell half-life to be 0.7 day, corresponding to a death rate *δ* = 1/day. The death rate of uninfected target cells is assumed to be 0.01/day, as previously estimated in patients for other viral infections^58^. The infection-free target cell concentration *T*_0_ is set at 10^6^/mL, such that at steady-state prior to infection yields *λ* = *dT*_0_ =10^4^/mL/day. The average eclipse period of an infected cell, 1/*k*, is set to 1 day, as previously estiamted^59^. The clearance rate of free CMV particles, *c*, is the same as estimated for other diverse viruses at 23/day (same as the rate estimated for HIV and HCV ^60,61^).

We now estimate the values of the other parameters in the model using the parameter values above and previously reported clinical data. First, Emery et al. estimated that the doubling time of the plasma CMV load to be 1 day in 18 bone marrow transplant recipients ^62^, which translates to an exponential growth rate of 0.69/day. This rate, *r*, corresponds to the dominant eigenvalue of the Jacobian matrix of Eqs. (1) at the infection free equilibrium. We find that *R*_0_ is related to *r* and other parameters by the equation

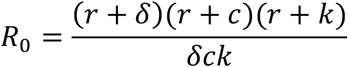

Substituting the values of *k*, *r, c* and *δ*, we get *R*_0_ =2.94.

From Eq. (1), we can calculate the set-point viral load (in the absence of treatment), *V*_0_:

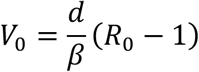

Emery et al. estimated that a typical set-point viral load is 10^5^ copies/mL in AIDS patients ^62^. Then, we calculate *β* as

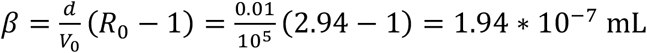

Finally, from Eq. 2, we calculate the value of *p* as

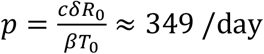

Note that the initial density of target cells *T*_0_ is not well established and that changes in the estimate of *T*_0_ will change the estimate of *p* without affecting the estimates of other parameters.

The table below summarizes the values of the parameters in the model:

| Parameters | Description | Value | Note |
| --- | --- | --- | --- |
| $\lambda$ | Influx rate of target cells | $10^4$ /mL/day | Set such that the target cell concentration without virus infection is $10^6$ copies/mL |
| $d$ | Death rate of target cells | 0.01 /day | Set to the death rate of T lymphocytes <sup>58</sup>. |
| $\beta$ | Virus infectivity | $1.94 * 10^{-7}$<br>mL | See calculation in the text |
| $k$ | Transition rate from eclipse phase to productive phase | 1 /day | Ref. <sup>59</sup> |
| $\delta$ | Death rate of infected cells | 1 /day | According to Ref. <sup>57</sup> |
| $p$ | Viral production rate | 349 /day | See calculation in the text |
| $c$ | Viral clearance rate | 23 /day | Set to the rate estimated for HIV and HCV <sup>60,61</sup>. |

### Calculation of probability of resistance mutants appearing in a host

The probably of resistant mutants appearing in a host is calculated using simulation of the ODE model (1) above. Viral load before initiation of therapy was assumed to be at steady-state, and therapy assumed to have 90% efficacy (i.e., *ε* = 0.9), starting at time 0. The model was then numerically solved and the total viral particles produced over time calculated from:

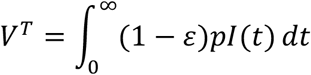

Simulation results of *V*(*t*), yield the total viral particles produced *V*^*T*^=7834 copies/mL. Note that this value is insensitive to the efficacy of the therapy as long as the therapy is effective and viral load declines exponentially under therapy^63^. For example, when *ε* = 0.7 or 0.99, we get the same number of particles produced, i.e. *V*^*T*^=7834 copies/mL. Assuming 15L of total body fluid in a person, this translated to approximately 7834*15*10^3^=1.1751*10^8^ viral particles produced during therapy. For GCV, where a single mutation is sufficient to generate a resistant mutant, the probability of generating one or multiple GCV resistant mutants based on a binomial draw from 10^8^ is 70%–100% (if *μ*=10^−3^ – 10^−4^ per replication, as reported^11^). For FD therapy, where the genetic barrier to resistance is high, the probability of generating resistance can be approximated as *V*^*T*^/*μ*^*n*^, where *n* is the number of mutations needed to generate resistance. Fig. 1C was generated assuming *μ* [10^−3^ – 10^−4^] and n=5, 10, 15, or 20.

### Double-stranded DNA-duplexes (preparation, modification, and nanoparticle construction)

FD^C^, FD^H^, FD^MCMV,^ FD^RhCMV^, Cy3- FD^C^ were made by annealing sequence specific oligonucleotides (see Extended Data Table 1 for sequences). Briefly, DNA oligonucleotides were obtained from Integrated DNA Technologies (San Jose, CA) and resuspended in annealing buffer (100mM Potassium acetate; 30mM HEPES, pH 7.5). Oligonucleotides of complementary sequences were mixed in equimolar amount, heated to 95°C for 2 minutes, and gradually cooled to 25°C over the period of 45 minutes in a S1000 Thermocycler (Bio-Rad) and stored at −20°C. Spherical nucleic acids (Cy3-FD^C^-SNA) were made from 3’ thiol labeled FD^C^ forward strand and 5’ cyanine 3- phosphoramidite (Cy3) tagged reverse strand oligos obtained from Bioneer, Inc (Alameda, CA) (see Extended Data Table 1 for sequences). Oligonucleotides were annealed as mentioned above and Cy3-FD-SNA were made as described previously^54^. Briefly, annealed duplexes were added to 10-nm citrate-stabilized gold nanoparticles (Sigma-Aldrich, St Louis, MO) (~3nmol oligonucleotide per 1ml of 10nM colloid). After 20 minutes, 10% SDS (Sigma-Aldrich, St Louis, MO) was added to bring down the concentration of SDS to 0.1% in phosphate buffer (0.01M, pH 7.4). 2.0M NaCl was then added to bring salt concentration to 0.1M. Duplexes were incubated under shaking conditions at room temperature for 30 min, followed by two successive additions of 2.0M NaCl 30 minutes apart to bring the final concentration of NaCl to 0.3M. The final mixture was incubated under shaking conditions overnight to complete the functionalization of duplexes on gold nanoparticles. The Cy3-FD-SNA was recovered by three centrifugation steps (13,000 rpm, 20 minutes), tested by dynamic light scattering to compare the diameter of gold nano particle (Extended Data Fig. 2b) before and after functionalizing with duplexes (Extended Data Fig. 2c). SNA were resuspended in 1x PBS buffer and stored at 4°C until used.

### Protein expression, EMSA and DNA binding assays

BL21 competent *E. Coli* cells (New England Biolabs Inc) were transformed with a pMALcXS plasmid encoding the C-terminus part of IE86 fused to the maltose binding protein, as described^64^, and induced with isopropyl b-D thiogalactoside (IPTG) in 1L luria broth (Thermofisher Scientific) containing ampicillin (Sigma-Aldrich). Purification for c-terminus IE86 protein: Cells were pelleted at 10,000 RPM for 30 minutes at 4°C and resuspended in 40mL lysis buffer (20mM HEPES, pH 7.4, 1M NaCl, 1mM EDTA, 1mM DTT, Roche protease inhibitor cocktail). Lysozyme was added at a final concentration of 1mg/ml and cells were incubated on ice for 30 minutes, followed by addition of 1mM PMSF and sonication on ice (sonicator at 40% amplitude, 10 seconds on, 30 seconds off-6 repeats). 1mM MgSO_4_, 0.1mg/ml DNAse (Sigma-Aldrich, St Louis, MO) and 25U/ml Benzonase (Sigma- Aldrich) were then added, cells further incubated on ice for 15-30 minutes, centrifuged at 12,000 rpm for 30 minutes at 4°C and the supernatant incubated with amylose resins (Sigma-Aldrich, St Louis, MO) for 2 hours at 4°C with gentle agitation. Batch bound resin was poured into a column and washed with 10 column volumes of lysis buffer, and eluted in elution buffer (20mM HEPES, pH 7.4°C, 250mM NaCl, 10mM Maltose, 1mM EDTA and 1mM DTT). The eluted protein was passed through a Superose 6 column (GE Healthcare Life Sciences) using gel filtration buffer (20mM HEPES, pH 7.4, 250mM NaCl and 1mM DTT) and used for EMSA or for binding assays. Cloning and purification of full length IE86 protein: Full length IE86 protein was codon optimized and cloned in pMALcx5 vector (New England Biolabs Inc.). Cells were transformed in BL-21 cells and IE86-FL-pMALcx5 transformed cells were grown in 1L luria broth (Thermofisher Scientific) containing ampicillin (Sigma-Aldrich) and induced for 1 hour with IPTG (Sigma Aldrich). Cells were pelleted and frozen in liquid nitrogen and stored at −80° freezer until used. Pellet was resuspended in 40mL lysis buffer (20mM HEPES, pH 7.4, 10% glycerol, 0.2M MgSO_4_, 1M NaCl, 1mM EDTA, 1mM DTT, Roche protease inhibitor cocktail). Lysozyme was added at a final concentration of 1mg/ml and cells were incubated on ice for 30 minutes, followed by addition of 1mM PMSF and sonication on ice (sonicator at 40% amplitude, 10 seconds on, 30 seconds off-6 repeats). 1mM MgSO_4_, 0.1mg/ml DNAse (Sigma-Aldrich, St Louis, MO) and 25U/ml Benzonase (Sigma- Aldrich) were then added, cells further incubated on ice for 15-30 minutes, centrifuged at 12,000 rpm for 30 minutes at 4°C and the supernatant incubated with amylose resins (Sigma-Aldrich, St Louis, MO) for 2 hours at 4°C with gentle agitation. Batch bound resin was poured into a column and washed with 10 column volumes of lysis buffer, and eluted in elution buffer (20mM HEPES, pH 7.4oC, 10% glycerol, 0.2M MgSO_4_, 250mM NaCl, 10mM Maltose, 1mM EDTA and 1mM DTT). The eluted protein was passed through a Superose 6 column (GE Healthcare Life Sciences) using gel filtration buffer (20mM HEPES, pH 7.4, 250mM NaCl 10% glycerol, 0.2M MgSO_4_, and 1mM DTT) and used for binding assays. Binding assays were performed by incubating duplexes of various lengths (as labeled in the figures) with purified MBP-IE86 or MBP-IE-FL for 30 minutes at room temperature and running on Superpose-6 column in the presence of gel filtration buffer and monitored oligomerized form of MBP-IE86 (at 13ml column volume on FPLC) in the presence of different sizes of FD^C^. To verify the sequence specific interaction between crs and MBP-IE86, EMSA was performed for crs and Δcrs oligonucleotides^65^. DNA oligonucleotides were obtained from Integrated DNA Technologies (San Jose, CA) (see Extended Data Table 1 for sequences) and resuspended in annealing buffer (100mM Potassium acetate; 30mM HEPES, pH 7.5). Oligonucleotides of complementary sequences were mixed in equimolar amount, heated to 95°C for 2 minutes, and gradually cooled to 25°C over 45 minutes in a S1000 Thermocycler (Bio-Rad) and stored at −20°C. Briefly, oligonucleotides were incubated with the c terminus-IE86 protein for 30 minutes in the presence of 1X binding buffer (10 mM HEPES-NaOH [pH 8.0], 50 mM KCl, 100 mM EDTA, and 5% glycerol, and subjected to electrophoresis on 1% agarose gel prepared in 1X TAE (Tris-acetate EDTA) buffer at 10V at 4°C. The gel was stained with ethidium bromide and imaged.

### Cell-culture conditions

ARPE-19 cells were maintained in a 1:1 mixture of Dulbecco’s Modified Eagle’s Medium (DMEM)/F-12 (Mediatech Inc.) with 10% fetal bovine serum (FBS) (HyClone) and 50U/ml Penicillin-Streptomycin (Mediatech Inc.) at 37oC and 5% CO_2_ in a humified incubator. MRC-5 fibroblasts, NIH 3T3 mouse fibroblast and Telo-RF were maintained in DMEM with 10% FBS and 50U/ml Penicillin and Streptomycin (Mediatech Inc.). Fomivirsen (NIH), Ganciclovir and Acyclovir (Sigma-Aldrich, St Louis, MO) were added to media at the indicated concentrations following virus inoculate removal. The ARPE-19 stably expressing the MIEP-IE86-GFP minimal circuit were previously described^22^. ARPE-19, Telo-RF (telomerase life extended rhesus fibroblast), NIH 3T3 mouse fibroblast and MRC-5 cell lines were obtained from ATCC.

### RNAseq analysis

Total RNA was isolated and purified using RNAeasy RNA extraction kit, Qiagen. RNA-seq libraries were prepared at the Gladstone Genomics Core with Ovation RNA-Seq System v2 kit (NuGEN). Total RNA (100ng) was reverse transcribed to synthesize the first-strand cDNA using a combination of random hexamers and a poly-T chimeric primer. RNA template was partially degraded by heating and the second strand cDNA was synthesized using DNA polymerase. The double-stranded DNA was amplified using single primer isothermal amplification (SPIA). (SPIA is a linear cDNA amplification process in which RNase H degrades RNA in DNA/RNA heteroduplex at the 5′-end of the double-stranded DNA, after which the SPIA primer binds to the cDNA and the polymerase starts replication at the 3′-end of the primer by displacement of the existing forward strand). Furthermore, random hexamers were used to amplify the second-strand cDNA linearly. The cDNA was fragmented with the Covaris S-series System according to the manufacturer’s recommendations to produce fragmented DNA with a fragment size in the desired range. Finally, libraries were generated (using seven cycles of amplification) from the SPIA amplified cDNA using the Ovation Ultralow Library System V2 (NuGEN). The RNA-seq libraries were analyzed by Bioanalyzer and quantified by qPCR (KAPA). High-throughput sequencing was done on one lane of the HiSeq 4000, single-read 50bp (Illumina). Completed libraries were sequenced on a NOVAseq 4000 at the UCSF Center for Advanced Technology. Sequencing reads were pseudo-aligned with kallisto^66^. Kallisto *count* function quantified transcript abundance for subsequent differential expression analysis with DESeq2^67^ in RStudio. All genes with less than 200 total reads across all samples were filtered out. The DESeq2 *results* function directly compared two sample types (i.e. 28 bp DNA vs Scramble), creating a table of genes with corresponding log2 fold change in expression, p-value and adjusted p-value. Genes with an adjusted p-value less than 0.05 and log2(fold change) either greater or less than 1 were extracted, generating lists of significantly up and down regulated genes. Gene lists were uploaded to DAVID Bioinformatics for Functional GO analysis to identify enriched biological processes^68^. Pathways were identified and ranked by fold enrichment. For the second set of analysis Up and down regulated gene lists were submitted to Gladstone Bioinformatics Core for additional analysis. For heatmap figures and functional enrichment analysis, the package DESeq2 computed normalized counts per million (CPM) mapped fragments from the raw count matrix using the robust median ratio method. Next, we log-transformed the CPM values (base 2) after adding a small number 0.01 to prevent logarithms of 0. This yielded a matrix of logCPM values for the four samples, which we subset for the genes with significant differential expression. We clustered the genes using *hopach* with cosine angle distance metric, other arguments specified as clusters=“best” and initord=“clust”^69^. This yielded nine clusters for the differential genes, whose logCPM values were plotted as a heatmap after setting the mean for each gene to zero. Then, we performed enrichment analyses to identify gene ontology (GO) terms for the clusters that had more than 100 genes with available Entrez IDs. For this, we used the enrichGO function from the clusterProfiler package to search for biological process sub-ontologies and assessed the significance of enrichment of a GO term relative to a background set of all the genes detected in our experiment^70^. We considered the terms with FDR-adjusted p < 0.05 as significantly enriched.

### Viral Replication Kinetics

The CMV TB40E-IE86-YFP virus was previously described^43^ and CMV AD169, GDGrK17, CMV GDGrP53, CMV 759rD100-1, CMV PFArD100 were obtained through the NIH AIDS Reagent Program. MCMV strain K181^71^ was kindly provided by Lewis Lanier (UCSF) and Rhesus CMV 68.1 EGFP virus^72^ was kindly provided by Peter Barry, UC-Davis. The clinical strain of HSV-1 17syn+ IE175-YFP^73^ was kindly provided by Roger Everett, MRC Virology Unit, Glasgow, Scotland that was passaged originally from a clinical isolate^74^. 1×10^6^ ARPE-19 cells were nucleofected with FD^C^, FD^H^ or FD^Scram^ using Amaxa nucleofector (Lonza). 24 hours post-nucleofection cells were infected in triplicate with 0.1 MOI of CMV (TB40E IE86-YFP) or HSV-1 (17syn+ IE175-YFP). Four days post infection, cells were harvested and subjected to three freeze-thaw cycles and used for 10-fold serial dilution titration on naïve ARPE-19 cells. As similar protocol was used to nucleofect MRC-5 cells prior to infection with AD169 and titration on naïve MRC-5 cells. Titration results were converted to PFU/ml using median tissue culture infectious dose (TCID50)^75^. Similarly, corneal tissues were dissociated using collagenase I (Sigma-Aldrich, St Louis, MO) for 30 minutes, subjected to three freeze-thaw cycles and used for TCID50 analysis.

### Quantitative PCR, flow cytometry and fluorescence microscopy

Genomic DNA from CMV or HSV-1 infected ARPE-19 cells, or mouse corneal tissue dissociated with collagenase I for 30 minutes, was extracted using DNeasy Blood and tissue kit (Qiagen). TLR9 expression analysis was performed by total cell RNA extraction using an RNEASY RNA isolation kit (74104; QIAGEN) and reverse transcription using a QuantiTet Reverse Transcription kit (205311; QIAGEN). Relative quantification of genomic DNA or cDNA was performed on a 7900HT Fast Real-Time PCR System (ThermoFisher Scientific, 4329003) using sequence specific primers (Extended Data Table 1) and Fast SYBR Green Master Mix (Applied Biosystems, 4385612). Flow cytometry was performed on BD FACS-Calibur (BD, Biosciences) at the Gladstone flow cytometry core and analyzed using FlowJo™ (FlowJo, LLC). Fluorescent microscopy of the cornea infected with HSV-1 was performed on a Zeiss Axiovert fluorescent microscope for DAPI (excitation at 345nm) and YFP (excitation at 514 nm). Images were analyzed using Image J software. Image quantification of YFP was based on a dual-thresholding for DAPI to generate image ‘masks’ (first a size thresholding for DAPI signal >10μm^2^ then an intensity threshold for DAPI intensity > 100 a.u.). YFP intensity was then quantified within the mask, for YFP particles > 2 μm^2^.

Imaging of CMV and HSV-1 cell entry was performed as described**. Briefly, HFFs and Vero cells grown on glass coverslips and infected with CMV TB40E IE86-YFP (MOI=5 imaged at 2h post-infection) or HSV-1 17syn+ IE175-YFP (MOI=20 imaged at 1h post-infection), respectively. Cells were fixed with 4% formaldehyde, permeabilized with 0.1% Triton-X100 for 15min and blocked in normal goat serum for 1h. HFFs were incubated with a mouse anti-pp65 primary antibody (clone 8F5, gift from the Shenk lab, 1:200 dilution) while Vero cells were incubated with a mouse anti-ICP5 primary antibody (Abcam ab6508, 1:200). HFFs and Vero cells were then incubated with a goat anti-mouse Alexa 594 secondary antibody (Invitrogen, A-11005, 1:250), mounted and imaged on a Zeiss Axiovert inverted fluorescence microscope (Carl Zeiss) with a 100X oil-immersion objective.

### Cell-death pathway inhibitor analysis

Analysis of cell-death pathways was performed using cell-death pathway inhibitors and staining for death-pathway markers. Briefly, cells were nucleofected with the FD^C^, FD^H^, or FD^Scram^ (25μM each) or mock nucleofected in the presence of one of the following inhibitors (auto= autophagy inhibitor, 3-methyladenine, authophagy inhibitor-M9281, Sigma Aldrich; apo= apoptosis inhibitor, NLRP3 inhibitor, CAS 256373-96-3, Calbiochem; fer= ferroptosis inhibitor, Ferrostatin-1, SML0583, Sigma Aldrich; nec=necroptosis inhibitor, Necrostatin-1, 4311-88-0, Sigma Aldrich) followed by infection with respective virus at MOI (0.1), at 48hpi, cells were trypsinized, stained with Zombie Aqua (BioLegend), and subjected to flow cytometry. Cells were stained for apoptosis marker (annexinV, Sigma Aldrich, cells harvested at 72hpi) and ferroptosis markers (cells were harvested at 24hpi) and subjected to flow cytometry.

### Corneal infection assays in mice

All experiments were performed with 6- to 10-week old male and female sibling Black 6 mice. Breeding pairs were purchased from Jackson laboratories (Bar Harbor, Maine) and maintained under pathogen-free conditions in the UCSF barrier facility. All animal experiments were conducted in accordance with procedures approved by the UCSF Institutional Animal Care and Use Committee. Corneal epithelial debridement was performed on mice as previously described^76^. Briefly, mice were anesthetized by isofluorane inhalation (Abbott Laboratories, Alameda, CA). The central part of the epithelium was removed down to the basement membrane using an Algerbrush II (Katena Products, Inc., Denville, NJ). 5μl of HSV-1 17syn+ YFP-IE175 (10^5^ pfu) were immediately applied to the debrided cornea. 6h later, mice were anesthetized again and 5μl of 25μM FD^H^, FD^Scram^ or PBS was applied to the cornea for 5 min. At the indicated time post infection, eyes were enucleated and the corneas were dissected to remove the lens, iris, and retina. Four incisions were made equal distances apart to aid in flattening the corneas. Fresh corneas were counterstained using 0.5μg/ml DAPI, mounted on slides with Fluoro-gel (Electron Microscopy Sciences, Hatfield, PA) and imaged on a Zeiss Axiovert Confocal microscope. Corneas were dissociated using collagenase I (Sigma-Aldrich, St Louis, MO), total DNA was extracted using DNeasy Blood & Tissue kit (QIAGEN), and subjected to qPCR using Fast SYBR green master mix (4385612; Applied Biosystems), analyzed on a 7900HT Fast Real-Time PCR System (4329003; Thermofisher Scientific). Virus titer was determined as described above.

### Statistical analysis

Statistical differences were determined by using the two-tailed unpaired Student’s *t* test and 2-way ANOVA followed by Tukey’s multiple comparisons test (GraphPad Prism, La Jolla, CA). A p-value less than 0.05 was considered statistically significant: *<0.05, **<0.01, ***<0.001, ****<0.0001, ns: not significant.

### Western blot analysis

10^6^ ARPE-19 cells were centrifuged at 2,000g for 10 min at 4°C and resuspended in PBS (Sigma-Aldrich, St Louis, MO). Cells were pelleted, resuspended in ice-cold RIPA lysis buffer (Sigma-Aldrich, St Louis, MO) by vortexing and incubated at 4°C for 30 min. The lysed samples were then pelleted at 13,000 rpm for 20 min at 4°C and supernatants were assayed by western blot as previously described^24^. Briefly, 20μg of cell lysate total protein was added to 1x loading buffer (100mM Tris-HCl (pH6.8), 200mM DTT, 4% SDS, 0.1% Bromophenol blue, 20% glycerol), and boiled for 10 minutes at 95°C. The lysed samples and precision plus kaleidoscope prestained protein marker (Bio-Rad) were then loaded on 10% Mini-PROTEAN TGX precast protein gel (Bio-Rad) in duplicate and ran at 90V for 2 hours in Tris-glycine running buffer (925mM Tris, 250mM Glycine and 0.1% SDS). One gel was stained for two hours at room temperature with coomassie blue stain (1g coomassie brilliant blue (Bio-Rad), Methanol (50%[v/v]), Glacial acetic acid (10%[v/v]), final volume to 1L with Milli-Q H_2_O, followed by destaining (40% methanol, 7% acetic acid in final volume of 1L in Milli-Q H_2_O. The gel was blotted on a PVDF membrane using a semi-dry transfer unit (Trans-Blot Semi-Dry Electrophotetic transfer cell, Bio-Rad) at 25V for 45 minutes. The membrane was blocked with Li-Cor Odyssey™ blocking buffer for 2 hours at room temperature with gentle agitation. IE86 protein was detected by a 2-hour incubation of the membrane with an anti-IE86 antibody (1:100 dilution, Mab 810, Chemicon) in Li-Cor Odyssey™ blocking buffer at room temperature followed by 3 5-minute wash steps (1x PBS + 0.01% Tween-20). The membrane was then incubated with a secondary Li-Cor detection antibody (1:20,000 dilution, goat anti-mouse 800CW) for 1 hour in the dark, washed three times in wash buffer (1x PBS + 0.01% Tween-20) and imaged on an Odyssey system (Li-Cor).

## References

1 Meylan, S., Andrews, I. W. & Collins, J. J. Targeting Antibiotic Tolerance, Pathogen by Pathogen. Cell 172, 1228–1238, doi:10.1016/j.cell.2018.01.037 (2018).

2 Lee, H. H., Molla, M. N., Cantor, C. R. & Collins, J. J. Bacterial charity work leads to population-wide resistance. Nature 467, 82–85, doi:10.1038/nature09354 (2010).

3 Goldberg, D. E., Siliciano, R. F. & Jacobs, W. R., Jr. Outwitting evolution: fighting drug-resistant TB, malaria, and HIV. Cell 148, 1271–1283, doi:10.1016/j.cell.2012.02.021S0092-8674(12)00221-8 [pii] (2012).

4 Piret, J. & Boivin, G. Antiviral drug resistance in herpesviruses other than cytomegalovirus. Rev Med Virol 24, 186–218, doi:10.1002/rmv.1787 (2014).

5 Lurain, N. S. & Chou, S. Antiviral drug resistance of human cytomegalovirus. Clin Microbiol Rev 23, 689–712, doi:10.1128/CMR.00009-1023/4/689 [pii] (2010).

6 Frobert, E. et al. Resistance of herpes simplex viruses to acyclovir: an update from a ten-year survey in France. Antiviral Res 111, 36–41, doi:10.1016/j.antiviral.2014.08.013 (2014).

7 Emery, V. C. & Griffiths, P. D. Prediction of cytomegalovirus load and resistance patterns after antiviral chemotherapy. Proc Natl Acad Sci U S A 97, 8039–8044, doi:10.1073/pnas.140123497 (2000).

8 Perelson, A. S. Modelling viral and immune system dynamics. Nat Rev Immunol 2, 28–36, doi:10.1038/nri700 (2002).

9 Coffin, J. M. HIV population dynamics in vivo: implications for genetic variation, pathogenesis, and therapy. Science 267, 483–489 (1995).

10 Cudini, J. et al. Human cytomegalovirus haplotype reconstruction reveals high diversity due to superinfection and evidence of within-host recombination. Proc Natl Acad Sci U S A 116, 5693–5698, doi:10.1073/pnas.1818130116[pii] (2019).

11 Drake, J. W. & Hwang, C. B. On the mutation rate of herpes simplex virus type 1. Genetics 170, 969–970, doi:10.1534/genetics.104.040410 (2005).

12 Renzette, N. et al. Limits and patterns of cytomegalovirus genomic diversity in humans. Proc Natl Acad Sci U S A 112, E4120–4128, doi:10.1073/pnas.1501880112[pii] (2015).

13 Renzette, N., Pfeifer, S. P., Matuszewski, S., Kowalik, T. F. & Jensen, J. D. On the Analysis of Intrahost and Interhost Viral Populations: Human Cytomegalovirus as a Case Study of Pitfalls and Expectations. J Virol 91, doi:e01976-16 [pii]10.1128/JVI.01976-16[pii] (2017).

14 Lu, Q., Hwang, Y. T. & Hwang, C. B. Mutation spectra of herpes simplex virus type 1 thymidine kinase mutants. J Virol 76, 5822–5828 (2002).

15 Lu, Q., Hwang, Y. T. & Hwang, C. B. Detection of mutations within the thymidine kinase gene of herpes simplex virus type 1 by denaturing gradient gel electrophoresis. J Virol Methods 99, 1–7 (2002).

16 Coen, D. M. & Schaffer, P. A. Two distinct loci confer resistance to acycloguanosine in herpes simplex virus type 1. Proc Natl Acad Sci U S A 77, 2265–2269 (1980).

17 Schnipper, L. E. & Crumpacker, C. S. Resistance of herpes simplex virus to acycloguanosine: role of viral thymidine kinase and DNA polymerase loci. Proc Natl Acad Sci U S A 77, 2270–2273, doi:10.1073/pnas.77.4.2270 (1980).

18 Goldner, T. et al. The novel anticytomegalovirus compound AIC246 (Letermovir) inhibits human cytomegalovirus replication through a specific antiviral mechanism that involves the viral terminase. J Virol 85, 10884–10893, doi:10.1128/JVI.05265-11 (2011).

19 Jaishankar, D. et al. An off-target effect of BX795 blocks herpes simplex virus type 1 infection of the eye. Sci Transl Med 10, doi:10.1126/scitranslmed.aan5861 (2018).

20 Cherrier, L. et al. Emergence of letermovir resistance in a lung transplant recipient with ganciclovir-resistant cytomegalovirus infection. Am J Transplant 18, 3060–3064, doi:10.1111/ajt.15135 (2018).

21 Jung, S., Michel, M., Stamminger, T. & Michel, D. Fast breakthrough of resistant cytomegalovirus during secondary letermovir prophylaxis in a hematopoietic stem cell transplant recipient. BMC Infect Dis 19, 388, doi:10.1186/s12879-019-4016-1 (2019).

22 Hochster, H. et al. Toxicity of combined ganciclovir and zidovudine for cytomegalovirus disease associated with AIDS. An AIDS Clinical Trials Group Study. Ann Intern Med 113, 111–117, doi:10.7326/0003-4819-113-2-111 (1990).

23 Troya, J. & Bascunana, J. Safety and Tolerability: Current Challenges to Antiretroviral Therapy for the Long-Term Management of HIV Infection. AIDS Rev 18, 127–137, doi:s113961211601 [pii] (2016).

24 Kumar, K. S. et al. Significant pulmonary toxicity associated with interferon and ribavirin therapy for hepatitis C. Am J Gastroenterol 97, 2432–2440, doi:10.1111/j.1572-0241.2002.05999.x (2002).

25 Bogdanove, A. J., Bohm, A., Miller, J. C., Morgan, R. D. & Stoddard, B. L. Engineering altered protein-DNA recognition specificity. Nucleic Acids Res 46, 4845–4871, doi:10.1093/nar/gky2894990010 [pii] (2018).

26 Pai, A. & Weinberger, L. S. Fate-Regulating Circuits in Viruses: From Discovery to New Therapy Targets. Annu Rev Virol 4, 469–490, doi:10.1146/annurev-virology-110615-035606 (2017).

27 Enquist, L. W. & Leib, D. A. Intrinsic and Innate Defenses of Neurons: Detente with the Herpesviruses. J Virol 91, doi:10.1128/JVI.01200-16 (2017).

28 Weller, S. K. & Coen, D. M. Herpes simplex viruses: mechanisms of DNA replication. Cold Spring Harb Perspect Biol 4, a013011, doi:10.1101/cshperspect.a013011 (2012).

29 Shenk, T. E. & Stinski, M. F. Human cytomegalovirus. Preface. Curr Top Microbiol Immunol 325, v (2008).

30 Mocarski, E. S., Shenk, T. & Pass, R. F. in Fields’ virology (ed David M. Knipe) 2708–2772 (Lippincott Williams & Wilkins, 2006).

31 Liu, B., Hermiston, T. W. & Stinski, M. F. A cis-acting element in the major immediate-early (IE) promoter of human cytomegalovirus is required for negative regulation by IE2. J Virol 65, 897–903 (1991).

32 Paterson, T. & Everett, R. D. The regions of the herpes simplex virus type 1 immediate early protein Vmw175 required for site specific DNA binding closely correspond to those involved in transcriptional regulation. Nucleic Acids Res 16, 11005–11025 (1988).

33 Teng, M. W. et al. An endogenous accelerator for viral gene expression confers a fitness advantage. Cell 151, 1569–1580, doi:10.1016/j.cell.2012.11.051 (2012).

34 Muller, M. T. Binding of the herpes simplex virus immediate-early gene product ICP4 to its own transcription start site. J Virol 61, 858–865 (1987).

35 O’Hare, P. & Hayward, G. S. Three trans-acting regulatory proteins of herpes simplex virus modulate immediate-early gene expression in a pathway involving positive and negative feedback regulation. J Virol 56, 723–733 (1985).

36 Isomura, H. et al. A cis element between the TATA Box and the transcription start site of the major immediate-early promoter of human cytomegalovirus determines efficiency of viral replication. J Virol 82, 849–858, doi:10.1128/JVI.01593-07 (2008).

37 Chaturvedi, S., Engel, R. & Weinberger, L. The HSV-1 ICP4 transcriptional auto-repression circuit functions as a transcriptional ‘accelerator’circuit. Frontiers in Cellular and Infection Microbiology 10, 265 (2020).

38 Hecker, M. & Wagner, A. H. Transcription factor decoy technology: A therapeutic update. Biochem Pharmacol 144, 29–34, doi:S0006-2952(17)30446-X [pii]10.1016/j.bcp.2017.06.122 (2017).

39 Werther, R. et al. Crystallographic analyses illustrate significant plasticity and efficient recoding of meganuclease target specificity. Nucleic Acids Res 45, 8621–8634, doi:10.1093/nar/gkx544 (2017).

40 Takeuchi, R., Choi, M. & Stoddard, B. L. Redesign of extensive protein-DNA interfaces of meganucleases using iterative cycles of in vitro compartmentalization. Proc Natl Acad Sci U S A 111, 4061–4066, doi:10.1073/pnas.1321030111[pii] (2014).

41 Seth, P. P., Tanowitz, M. & Bennett, C. F. Selective tissue targeting of synthetic nucleic acid drugs. J Clin Invest 129, 915–925, doi:10.1172/JCI125228 (2019).

42 Stein, C. A. & Castanotto, D. FDA-Approved Oligonucleotide Therapies in 2017. Mol Ther 25, 1069–1075, doi:S1525-0016(17)30122-3 [pii]10.1016/j.ymthe.2017.03.023 (2017).

43 Vardi, N., Chaturvedi, S. & Weinberger, L. S. Feedback-mediated signal conversion promotes viral fitness. Proc Natl Acad Sci U S A 115, E8803–E8810, doi:10.1073/pnas.1802905115 (2018).

44 Hong, S. H. et al. Molecular crosstalk between ferroptosis and apoptosis: emerging role of ER stress-induced p53-independent PUMA expression. Oncotarget 8, 115164–115178, doi:10.18632/oncotarget.23046[pii] (2017).

45 Everett, R. D., Murray, J., Orr, A. & Preston, C. M. Herpes simplex virus type 1 genomes are associated with ND10 nuclear substructures in quiescently infected human fibroblasts. J Virol 81, 10991–11004, doi:10.1128/JVI.00705-07 (2007).

46 Luecke, S. et al. cGAS is activated by DNA in a length-dependent manner. EMBO Rep 18, 1707–1715, doi:10.15252/embr.201744017 (2017).

47 Mulamba, G. B., Hu, A., Azad, R. F., Anderson, K. P. & Coen, D. M. Human cytomegalovirus mutant with sequence-dependent resistance to the phosphorothioate oligonucleotide fomivirsen (ISIS 2922). Antimicrob Agents Chemother 42, 971–973 (1998).

48 Cunningham, E. T. Cytomegalovirus: ophthalmic perspectives on a pervasive pathogen. Expert Review of Ophthalmology 6, 489–491, doi:10.1586/eop.11.50 (2011).

49 Elia, M. H., J.J., and Gaudio, P.A. in EyeNet Magazine 37–38 (2016).

50 Lahmidi, S., Yousefi, M., Dridi, S., Duplay, P. & Pearson, A. Dok-1 and Dok-2 Are Required To Maintain Herpes Simplex Virus 1-Specific CD8(+) T Cells in a Murine Model of Ocular Infection. J Virol 91, doi:10.1128/JVI.02297-16 (2017).

51 Adams, D. et al. Patisiran, an RNAi Therapeutic, for Hereditary Transthyretin Amyloidosis. N Engl J Med 379, 11–21, doi:10.1056/NEJMoa1716153 (2018).

52 Kanasty, R., Dorkin, J. R., Vegas, A. & Anderson, D. Delivery materials for siRNA therapeutics. Nat Mater 12, 967–977, doi:10.1038/nmat3765 (2013).

53 Khvorova, A. & Watts, J. K. The chemical evolution of oligonucleotide therapies of clinical utility. Nat Biotechnol 35, 238–248, doi:10.1038/nbt.3765 (2017).

54 Rosi, N. L. et al. Oligonucleotide-modified gold nanoparticles for intracellular gene regulation. Science 312, 1027–1030, doi:10.1126/science.1125559 (2006).

55 Asberg, A. et al. Lessons Learned From a Randomized Study of Oral Valganciclovir Versus Parenteral Ganciclovir Treatment of Cytomegalovirus Disease in Solid Organ Transplant Recipients: The VICTOR Trial. Clin Infect Dis 62, 1154–1160, doi:10.1093/cid/ciw084 (2016).

56 Diekmann, O., Heesterbeek, J. A. & Roberts, M. G. The construction of next-generation matrices for compartmental epidemic models. J R Soc Interface 7, 873–885, doi:10.1098/rsif.2009.0386 (2010).

57 Rose, J. et al. Novel decay dynamics revealed for virus-mediated drug activation in cytomegalovirus infection. PLoS Pathog 13, e1006299, doi:10.1371/journal.ppat.1006299 (2017).

58 Mohri, H., Bonhoeffer, S., Monard, S., Perelson, A. S. & Ho, D. D. Rapid turnover of T lymphocytes in SIV-infected rhesus macaques. Science 279, 1223–1227, doi:10.1126/science.279.5354.1223 (1998).

59 Emery, V. C., Hassan-Walker, A. F., Burroughs, A. K. & Griffiths, P. D. Human cytomegalovirus (HCMV) replication dynamics in HCMV-naive and -experienced immunocompromised hosts. J Infect Dis 185, 1723–1728, doi:10.1086/340653 (2002).

60 Ramratnam, B. et al. Rapid production and clearance of HIV-1 and hepatitis C virus assessed by large volume plasma apheresis. Lancet 354, 1782–1785, doi:10.1016/S0140-6736(99)02035-8 (1999).

61 Guedj, J. et al. Modeling shows that the NS5A inhibitor daclatasvir has two modes of action and yields a shorter estimate of the hepatitis C virus half-life. Proc Natl Acad Sci U S A 110, 3991–3996, doi:10.1073/pnas.1203110110 (2013).

62 Emery, V. C., Cope, A. V., Bowen, E. F., Gor, D. & Griffiths, P. D. The dynamics of human cytomegalovirus replication in vivo. J Exp Med 190, 177–182, doi:10.1084/jem.190.2.177 (1999).

63 Bonhoeffer, S., May, R. M., Shaw, G. M. & Nowak, M. A. Virus dynamics and drug therapy. Proc Natl Acad Sci U S A 94, 6971–6976, doi:10.1073/pnas.94.13.6971 (1997).

64 Macias, M. P. & Stinski, M. F. An in vitro system for human cytomegalovirus immediate early 2 protein (IE2)-mediated site-dependent repression of transcription and direct binding of IE2 to the major immediate early promoter. Proc Natl Acad Sci U S A 90, 707–711, doi:10.1073/pnas.90.2.707 (1993).

65 Asmar, J., Wiebusch, L., Truss, M. & Hagemeier, C. The putative zinc finger of the human cytomegalovirus IE2 86-kilodalton protein is dispensable for DNA binding and autorepression, thereby demarcating a concise core domain in the C terminus of the protein. J Virol 78, 11853–11864, doi:10.1128/JVI.78.21.11853-11864.2004 (2004).

66 Bray, N. L., Pimentel, H., Melsted, P. & Pachter, L. Near-optimal probabilistic RNA-seq quantification. Nat Biotechnol 34, 525–527, doi:10.1038/nbt.3519 (2016).

67 Love, M. I., Huber, W. & Anders, S. Moderated estimation of fold change and dispersion for RNA-seq data with DESeq2. Genome Biol 15, 550, doi:10.1186/s13059-014-0550-8 (2014).

68 Huang da, W., Sherman, B. T. & Lempicki, R. A. Bioinformatics enrichment tools: paths toward the comprehensive functional analysis of large gene lists. Nucleic Acids Res 37, 1–13, doi:10.1093/nar/gkn923 (2009).

69 Pollard, K. S. & van der Laan, M. J. Statistical inference for simultaneous clustering of gene expression data. Math Biosci 176, 99–121, doi:10.1016/s0025-5564(01)00116-x (2002).

70 Yu, G., Wang, L. G., Han, Y. & He, Q. Y. clusterProfiler: an R package for comparing biological themes among gene clusters. OMICS 16, 284–287, doi:10.1089/omi.2011.0118 (2012).

71 Morley, P. J., Ertl, P. & Sweet, C. Immunisation of Balb/c mice with severely attenuated murine cytomegalovirus mutants induces protective cellular and humoral immunity. J Med Virol 67, 187–199, doi:10.1002/jmv.2207 (2002).

72 Chang, W. L. & Barry, P. A. Cloning of the full-length rhesus cytomegalovirus genome as an infectious and self-excisable bacterial artificial chromosome for analysis of viral pathogenesis. J Virol 77, 5073–5083 (2003).

73 Everett, R. D., Sourvinos, G. & Orr, A. Recruitment of herpes simplex virus type 1 transcriptional regulatory protein ICP4 into foci juxtaposed to ND10 in live, infected cells. J Virol 77, 3680–3689 (2003).

74 Brown, S. M., Ritchie, D. A. & Subak-Sharpe, J. H. Genetic studies with herpes simplex virus type 1. The isolation of temperature-sensitive mutants, their arrangement into complementation groups and recombination analysis leading to a linkage map. J Gen Virol 18, 329–346, doi:10.1099/0022-1317-18-3-329 (1973).

75 Reed, L. J. M., H. A Simple method of estimating fifty percent endpoints. Journal of Epidemiology 27, 493–497 (1938).

76 Chan, M. F. & Werb, Z. Animal Models of Corneal Injury. Bio Protoc 5, e1516 (2015).

77 Skouboe MK, Knudsen A, Reinert LS, et al. STING agonists enable antiviral cross-talk between human cells and confer protection against genital herpes in mice. PLoS Pathog. 2018;14(4):e1006976 (2018).

