## Extended data figures 1-9 and Table 1 for "Transcriptional Feedback Disruption Yields Escape-Resistant Antivirals"

**<u>Extended Data Figure 1</u>: Simulations and in-vitro analyses show crs-encoding DNA duplexes could competitively homo-oligomerize IE86 disrupting IE86 negative feedback and increasing IE86 levels.** (a) Numerical simulations of an experimentally validated Ordinary Differential Equation (ODE) model of the Major Immediate Early circuit of CMV[33, 36] modified to include FDs as described in Methods for four different values of parameter $k_1$ [0.01, 0.1, 1, 10]. All other parameters are kept constant and initial levels of FD DNA duplex (t=0) are varied according to color legend (at right). (b) Electrophoresis mobility shift assays (EMSA) verifying that the c-terminus of IE86 specifically interacts with crs-containing DNA duplexes. Digoxygenin (DIG)-labeled DNA-duplex probes of the crs and Δcrs sequences, as previously described[65], were used for binding and detection. EMSA was performed after incubating DIG-labeled DNA probes with increasing concentrations of IE86 protein (0μM to 20μM) for 30 minutes at room temperature (lanes 1–6). To determine if the IE86- crs interaction was sequence specific, either 20-fold of unlabeled crs DNA duplex (*) was added (lane 7) or 20-fold unlabeled Δcrs DNA duplex (**) was added (lanes 7 and 10). (c) crs-encoding DNA duplexes ranging from 20–30bp can efficiently catalyze IE86 homo-multimer formation. Size exclusion chromatography of mixtures of DNA duplexes of different lengths (as indicated) with c-terminus of IE86 protein. The IE86 homomultimer protein-DNA complex elutes at ~13ml column volume (red bar). (d) Size-exclusion chromatography of Maltose Binding Protein (MBP) together with $FD^C$ indicates that MBP does not oligomerize in the presence of $FD^C$ excluding an MBP-mediated protein-DNA interaction underlying complex formation and oligomerization. (e) Delivery and stability of $FD^C$ (28bp-DNA duplex) into cells; fluorescence micrographs of ARPE-19 cells four days post nucleofection with cy3-tagged $FD^C$ (left) or free cy3 dye (right). (f) Phosphorothioation enhances efficacy of $FD^C$: flow-cytometry analysis of IE86 feedback-reporter cell four days after nucleofection with either 25μM of 28-bp scramble DNA duplex, $FD^C$ (four phosphorothioate bonds), or a non-phosphorothioated version of $FD^C$. (g) Flow cytometry dot plots of the IE86 negative-feedback reporter cell line after nucleofection with increasing doses of the $FD^C$ and 100μM of the $FD^{Scram}$ (negative control). % GFP+ population was quantified two days post nucleofection (SSC = side scatter). Sequence schematics of the unmodified DNA duplex $FD^C$(N), the phosphorothioated $FD^C$ (* indicates phosphorothioated bases), and 28bp scrambled DNA (also referred to as $FD^{Scram}$), which also contains phosphorothioated bases. (h) Top: Quantitative (LICOR™) Western blot analysis of IE86 from cell lysates of panel A at two days post FD nucleofection; Bottom: Coomassie blue loading control for western blot. (i) Detection of apoptosis by TUNEL assay in IE86 negative-feedback reporter cells. Cells were nucleofected with 25μM $FD^C$, $FD^{Scram}$, or mock nucleofected, harvested by trypsinization 72 hours later, stained with TUNEL marker (Biotium inc.) and analyzed by flow cytometry.

**<u>Extended Data Figure 2</u>: $FD^C$ catalyzes oligomerization of full-length IE86 protein, nanoparticles increase FD cellular uptake, and FD concatemers enhance IE86 sequestration and feedback disruption.** (a) Size-exclusion chromatography of purified full-length IE86 protein shows that $FD^C$ efficiently catalyzes IE86 oligomerization, eluting at ~11ml column volume. (b, c) Dynamic light scattering measurements to determine diameters of gold nanoparticle before (b) and after (c) addition of 28bp $FD^C$ DNA duplexes to generate spherical nucleic acid (SNA) nanoparticles. Cy3-tagged $FD^C$ conjugated to 10-nm gold nanoparticles to generate SNAs. (d) Cellular uptake of $FD^C$ SNA nanoparticles is more efficient than uptake of 'free' $FD^C$ DNA duplexes. Cells were incubated in culture with the $FD^C$ SNAs, 4 days later cells were assayed for Cy3 by fluorescent microscopy. Micrographs of ARPE-19 cells at 4 days post incubation with cy3-tagged $FD^C$-SNA, cy3-tagged 'free' $FD^C$, or unlabeled 10-nm gold nanoparticles lacking cy3 (as a control) are shown in the cy3 fluorescence channel (left) and cy3 overlap with bright field channel (right). (e) Size-exclusion chromatography of purified $IE86_C$ protein fragment (n-terminus tagged with MBP) incubated with $FD^C$ containing one crs sequence (FD 1x) of two concatenated crs sequences (FD 2X-concat) for 30 minutes at room temperature (see Extended Data Table 1 for FD sequences). Oligomerized fraction (% absorbance at ~13ml fraction at OD280) was compared for both the samples. (f) Flow cytometry analysis of the IE86 feedback circuit reporter cell line two days after nucleofection with 25μM FD (1X), FD (2X)-concat, or the scrambled DNA oligo ($FD^{Scram}$, negative control). (g) Flow cytometry analysis of the IE86 feedback circuit reporter cell line two days after nucleofection with 25μM $ssFD^C$, or the scrambled DNA oligo ($FD^{Scram}$).

__Extended Data Figure 3__. **RNAseq analysis indicates that disrupting IE86 negative feedback upregulates apoptosis in the feedback-reporter cell line.** (a) Heatmap showing the logCPM (log counts per million) values for clusters of differentially expressed genes from RNAseq of cells nucleofected with either $FD^C$ or $FD^{Scram}$. (biological duplicates shown for each condition). The clusters are numbered on the left side of the plot and the sample IDs are indicated at the bottom. (b) Differentially expressed genes clustered by the logCPM values of FD and $FD^{Scram}$ samples [see methods]. Cluster 4 is a set of upregulated genes with enrichment of GO terms related to cell migration, endocytosis, and apoptosis. The y-axis shows the enriched terms and the x-axis shows GeneRatio, (the fraction of genes associated with a term that were among the genes in this cluster). The sizes of the data point represent the number of genes in the cluster associated with the respective term and the colors represent the FDR-adjusted p-values. The apoptotic signaling pathway term is high in GeneRatio and Gene Count with an adjusted p-value < 0.05. (c) The logCPM values of clusters 4 and 5 plotted as a heatmap after setting the mean for each gene to zero sorted by their logFC. Genes associated with apoptosis terms that had unadjusted p < 0.01 are labeled.

__Extended Data Figure 4__. **FD DNA duplexes do not prevent virus entry into cells.** (a) Flow cytometry dot plots of naïve or $FD^C$-nucleofected ARPE-19 cells after infection with CMV TB40E-IE86-YFP (MOI 0.1). Cells were infected 24h after nucleofection with $FD^C$ and analyzed at 4 hpi (hours post infection). (b) Flow cytometry dot plots of naïve or $FD^H$-nucleofected ARPE-19 cells after infection with HSV-1 strain 17syn+ IE175-YFP (MOI 0.1). Cells were infected 24h after nucleofection with $FD^H$ and analyzed at 4 hpi. (c) Left: Fluorescent micrographs of HFF cells (nucleofected +/– $FD^C$) then infected with CMV TB40E-IE86-YFP, imaged at 2 hpi (MOI=5), and stained for pp65 protein (red, 594nm) and DAPI (blue, 405nm). Right: Fluorescent micrograph of Vero cells (nucleofected +/– $FD^H$) then infected with HSV-1 17syn+ IE175-YFP and imaged at 1 hpi (MOI=20) and stained for ICP5 protein (red, 594nm) and DAPI (blue, 405nm).

__Extended Data Figure 5__: **Feedback disruption inhibits viral replication in a broad range of species-specific herpesviruses (including drug-resistant strains); FDs diffuse into naïve mouse corneal cells and increase IE175 expression following HSV-1 infection of mouse corneas.** (a) Sequence homology for the crs of human CMV, murine CMV (MCMV). and rhesus CMV (RhCMV), green represents sequence homology whereas red represents divergence. (b) $FD^{MCMV}$ interferes with MCMV replication. NIH

3T3 mouse fibroblast cells were nucleofected with either 25μM $FD^{MCMV}$, $FD^{Scram}$, or mock nucleofected 24 hours prior to MCMV infection at MOI = 0.1. 4 dpi (days post infection), virus titers were assayed by TCID50 on 3T3 cells. (c) $FD^{RhCMV}$ interferes with RhCMV replication. Fluorescent micrographs, at 4-day post nucleofection of Telo-RF cells with either 25μM $FD^{RhCMV}$ or ($FD^{Scram}$/mock), cells were infected with RhCMV (RhCMV 68.1 GFP) at MOI = 0.1. Virus titers assayed by TCID-50 at 4 dpi. (d) $FD^{C}$ interferes with replication of ganciclovir-resistant (GCVR) and foscarnet-resistant (FOSR) CMV strains. MRC-5 cells were nucleofected with 25μM $FD^{C}$ or $FD^{Scram}$ and 24 hours later were infected with either parent CMV AD169 (control) or GCVR or FOSR strains (CMV GDGrK17, CMV GDGrP53, CMV 759rD100-1, CMV PFArD100) at MOI = 0.1. Virus titers were assayed by TCID-50 at 4 dpi. (e) $FD^{H}$ interferes with HSV-1 (KOS strain) replication. Vero cells were nucleofected with either 25μM $FD^{H}$ or ($FD^{Scram}$/mock), and at 24 hours post nucleofection, cells were infected with HSV-1(KOS strain) at MOI = 0.1, virus was titered at 4 dpi by TCID50. (f) Fluorescence micrographs of dissected mouse corneas incubated with 25μM Cy3-tagged $FD^{H}$ for 1 hour. Corneas were washed three times in PBS and immediately imaged. (g) YFP and DAPI fluorescence micrographs of dissected mouse corneas +/- $FD^{H}$ at one day post HSV-1 infection (17syn+ IE175-YFP). (h) Triplicate repeats of CMV-infected ARPE-19 (left) and HSV-1 infected Vero (right) cells stained with cell death markers in the presence of FDs. (left) ARPE-19 cells were nucleofected with 25μM $FD^{C}$ (or mock/$FD^{Scram}$), 24 hours later cells were infected with CMV (TB40/E) IE86-YFP virus (MOI=1), and at 48 h post infection stained with Annexin V. (right) Vero were nucleofected with 25μM $FD^{H}$ (or mock/$FD^{Scram}$), at 24 hour post nucleofection, cells were infected with HSV-1 (17syn+ strain) IE175-YFP (MOI=1), and at 24 hour post infection cells were harvested and stained with a ferroptosis marker.

**Extended Data Figure 6: Feedback disruptors induce cell death only in presence of virus infection via specific cell-death pathways as identified by small-molecule inhibitors (cell-death rescue assay).** (a) Representative flow cytometry dot plot of live-dead analysis using Zombie Aqua of uninfected (naïve) ARPE-19 cells (i.e., control) showing gating for dead cells. Dead cell gate drawn was drawn by comparing uninfected ARPE-19 in panel a to CMV-infected ARPE-19 cells in panel e (left-most column). Percentage of dead ARPE-19 cells in uninfected culture is < 1%. (b) uninfected ARPE-19 cells nucleofected with $FD^{Scram}$ or $FD^{C}$; nucleofection of $FD^{C}$ does not generate significant increases in dead cells compared to nucleofection of $FD^{Scram}$ control (mean of biological triplicates shown). (c) Flow cytometry live-dead analysis using Zombie Aqua of uninfected (naïve) Vero cells (i.e., control) showing gating for dead cells. Dead cell gate was drawn by comparing uninfected Vero in panel d to HSV-1-infected Vero cells in panel f (left-most column). Percentage of dead cells is < 1%. (d) uninfected Vero cells nucleofected with $FD^{Scram}$ or $FD^{H}$ (mean of biological triplicates shown); nucleofection of $FD^{H}$ does not generate significant increases in dead cells compared to nucleofection of $FD^{Scram}$ control. (e) ARPE-19 cells infected with TB40E-IE86-YFP virus (24 hours post nucleofection of $FD^{C}$ or $FD^{Scram}$) in the presence of the indicated cell-death inhibitors (auto = authophagy inhibitor; apo = apoptosis inhibitor; nec = necroptosis inhibitor; fer = ferroptosis inhibitor). Cells were harvested at 24 hpi, stained for dead cells with Zombie Aqua (BioLegend), and analyzed by flow cytometry. The experiment performed for three biological replicates. (f) Vero cells nucleofected with $FD^{Scram}$ or $FD^{H}$ were infected with HSV-1 17syn+ IE175-YFP virus (24 hours post nucleofection) in the presence of the indicated cell-death inhibitors (as in panel d). Cells were harvested at 24 hpi, stained for dead cells with Zombie Aqua (BioLegend), and analyzed by flow cytometry. The experiment was performed in three biological replicates.

<u>Extended Data Figure 7</u>: **Feedback disruption does not induce apoptosis in uninfected bystander cells; FDs do not act through innate immune signalling; ACV and fomivirsen rapidly select for resistance; nanoparticle FD efficacy**. (a) Schematic of the experiment to examine if feedback disruption in infected cells affects viability of uninfected neighbouring 'bystander' cells. ARPE-19 cells were nucleofected with 25μM $FD^C$ and 24 hours post nucleofection cells were infected with CMV TB40E-IE86-YFP (MOI = 0.5). At 2 hpi, cells were washed twice in PBS to remove any attached virus, and co-cultured with naïve mCherry-expressing ARPE-19 cells ('bystanders'). Cells were analysed by flow cytometry at 36 hpi (i.e., once FD-mediated cytotoxicity was present but prior to virus release). (b) Flow cytometry analysis of YFP (IE86 expression), mCherry (bystander cells), and AnnexinV (apoptosis) in co-cultures at 36 hpi. (c) AnnexinV staining in the IE86-YFP positive (infected) cells versus mCherry-positive (bystander) cells from the co-culture at 36 hpi. Bystander cells do not exhibit apoptosis whereas feedback disrupted CMV-infected cells do. (d) FDs do not activate the TLR9 response. qPCR analysis of TLR9 expression in ARPE-19 cells 4 days after nucleofection of either a TLR9-activating oligonucleotide (ODN2216)[77], 25μM $FD^C$, or mock nucleofected. Total RNA was extracted from cells and corresponding cDNA was quantified by qPCR using sequence-specific primers for TLR9 (see Extended Data Table 1). (e) FDs do not act through the cGAS-STING pathway. ARPE-19 (low cGAS-STING activity) and MRC-5 cells (high cGAS-STING activity) were nucleofected with 25μM $FD^C$ or mock and infected with CMV TB40E-IE86-YFP or AD169 (MOI=0.1), respectively. Virus titers were assayed by TCID-50 at 4 dpi. (f) $FD^C$ decreases CMV titers below detection by day 60 in continuous culture (see Fig 3e). (g) Viral titers from continuous cultures in the presence/absence of 25μM Fomivirsen. Mock-nucleofected or Fomivirsen-nucleofected ARPE-19 cells were infected with CMV TB40E-IE86-YFP (MOI=0.1); at 4 dpi, supernatants were transferred to infect naïve ARPE-19 cells (+/- Fomivirsen, no dose escalation used) and transfers repeated every 4 days until day 44 (see schematic of continuous culture, Fig 3D). (h) Viral titers from continuous cultures (rounds 3 to 5), +/- 25μM Fomivirsen (no dose escalation used). A positive slope in the titers in Fomivirsen samples (i.e., outgrowth of fomivirsen-resistant virus) is observed beginning at round 3 of infection. (i) HSV-1 resistance to acyclovir (ACV). Viral titers of ARPE-19 cells infected with HSV-1 (strain 17syn+ IE175-YFP; MOI=1) +/- 100μM ACV (no dose escalation used) and supernatant transferred every 2 days over 3 consecutive rounds of infection. Virus titers were assayed by TCID50 every 2 days post transfer; positive slope in the titers (i.e., outgrowth of acv-resistant virus) is evident despite 100μM ACV. (j) Flow cytometry analysis of IE86-GFP feedback reporter cell line treated with 10-nm gold nanoparticles (control) or 10-nm $FD^C$-SNAs at 4 days post incubation. (k) ARPE-19 cells were treated with gold nanoparticles or $FD^C$ SNAs and subsequently infected with CMV followed by viral titering by TCID50 at 4 dpi (see Extended Data Table 1 for sequences).

<u>Extended Data Figure 8</u>: **The CMV Major Immediate Early promoter-enhancer DNA-protein interaction domains exhibit no dominant (i.e., population sweep) mutations over 60 days of $FD^C$ treatment.** Sequencing analysis of the CMV promoter region from the continuous culture experiment in the presence of $FD^C$ (Fig. 3e). Viral DNA was extracted and 250bp upstream and downstream of the crs was amplified by PCR (see Extended Data Table 1 for primer sequences), Sanger sequenced, and then aligned with

the viral promoter sequence for homology analysis. No acquired consensus mutations are observed over 14 replication cycles.

**Extended Data Figure 9: The IE86 protein-protein and DNA-protein interaction domains (exon 5) exhibits no dominant (i.e., population sweep) mutations over 60 days of $FD^C$ treatment.** Extended $FD^C$ treatment over 60 days does not appear to generate dominant mutations. Sanger sequencing of IE86 (Exon 5) from virus in the continuous culture experiment in the presence of $FD^C$ (Fig. 3e). Viral DNA was extracted, IE86 exon 5 was PCR amplified (see Extended Data Table 1 for primer sequences), Sanger sequenced, and aligned with the IE86 sequence for homology analysis. Despite apparent transient SNPs (e.g., at nucleotide position 640), no mutations 'sweep' the population or are maintained in the consensus sequence over time.

### Extended Data Figure 1 (Chaturvedi et al.)

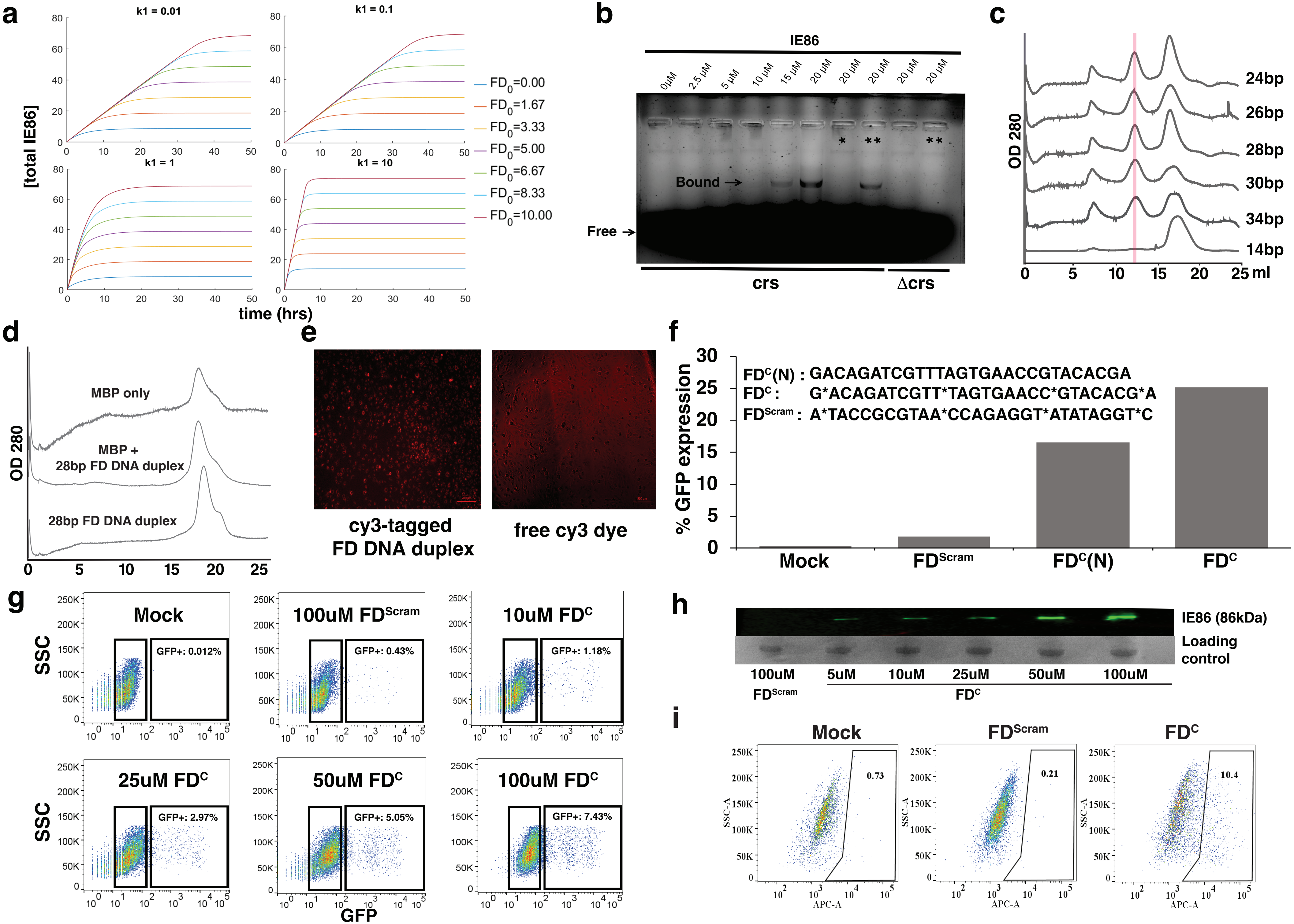

### Extended Data Figure 2 (Chaturvedi et al.)

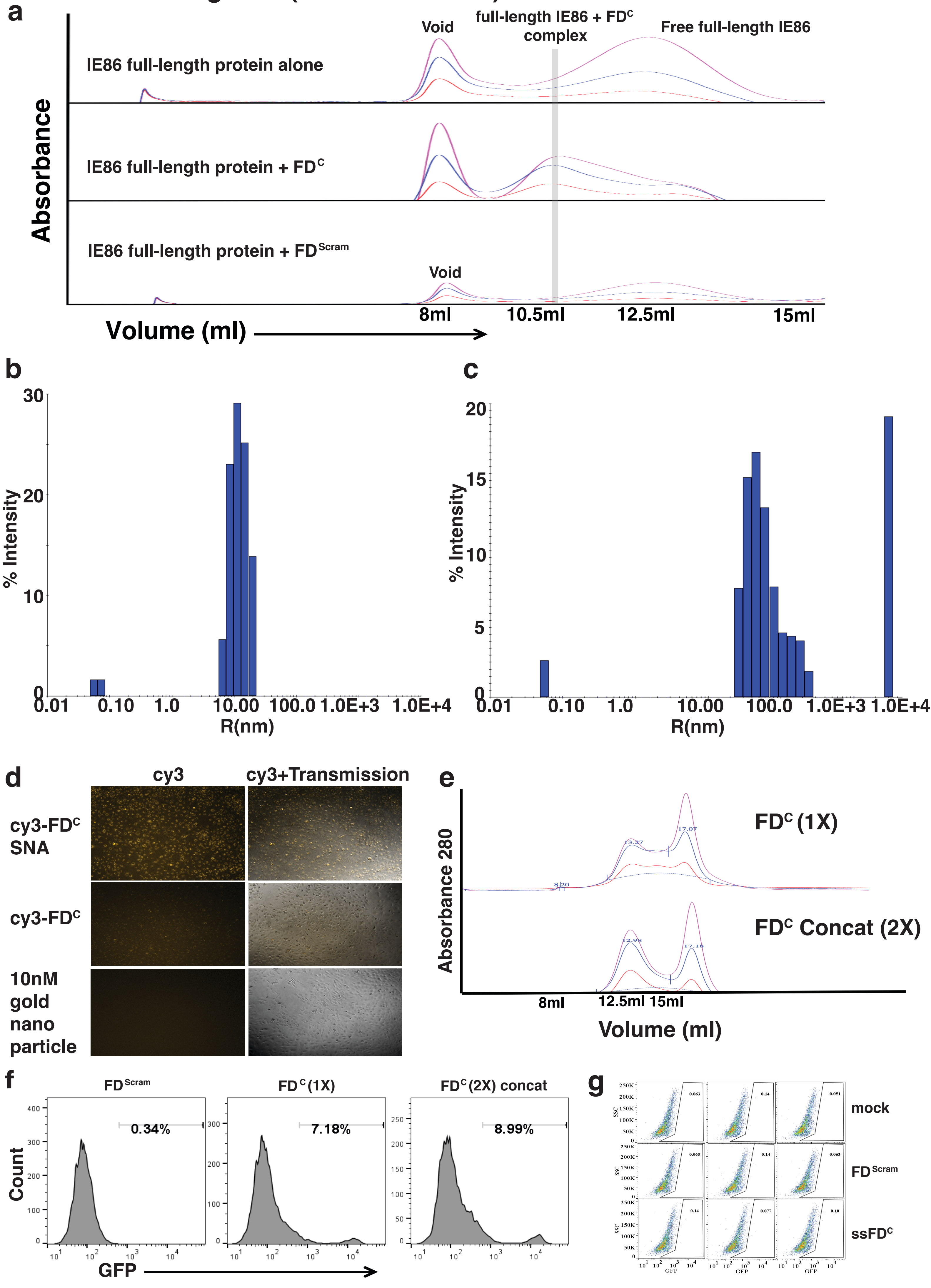

### Extended Data Figure 3 (Chaturvedi et al.)

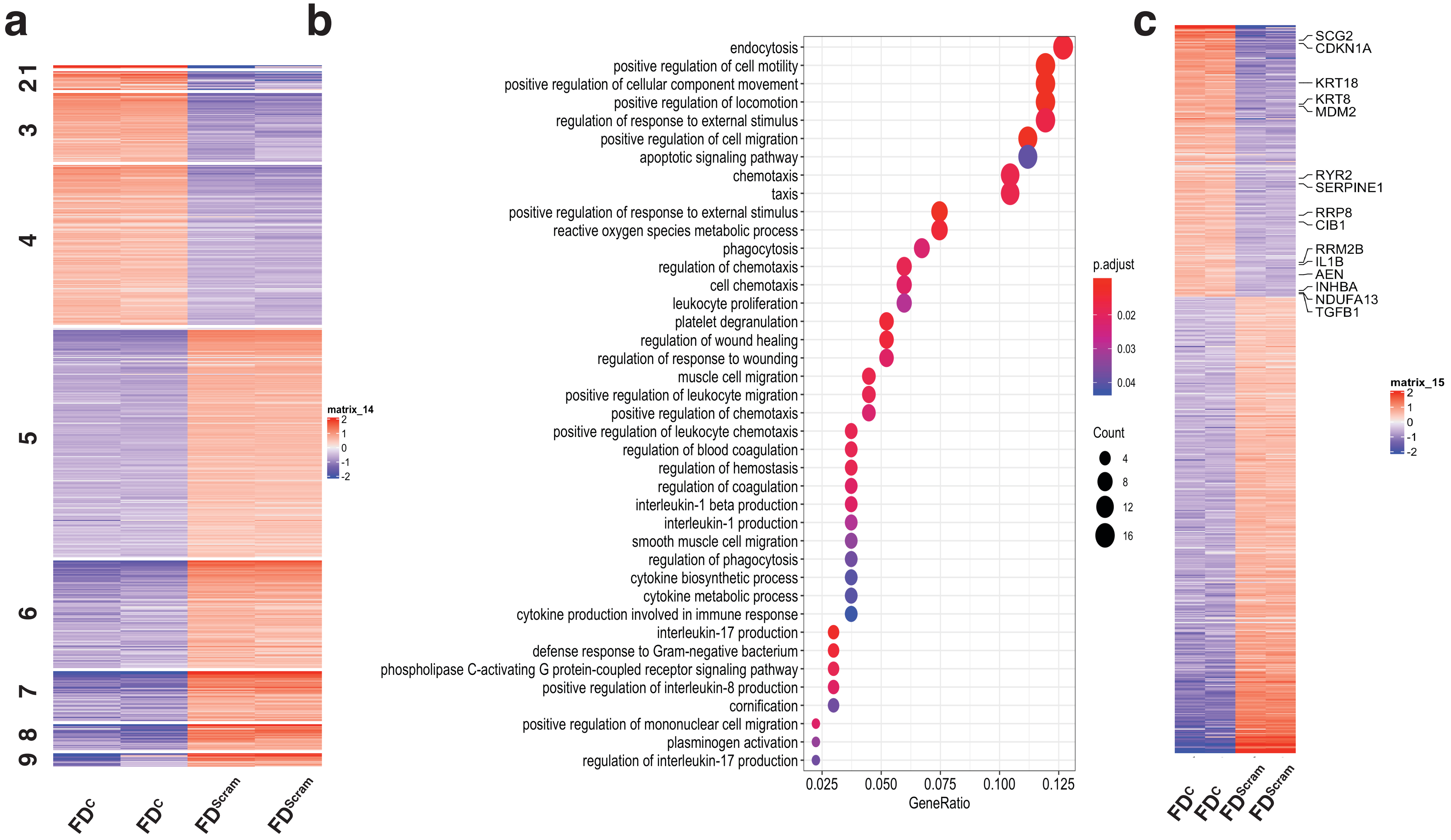

Extended Data Figure 4 (Chaturvedi et al.)

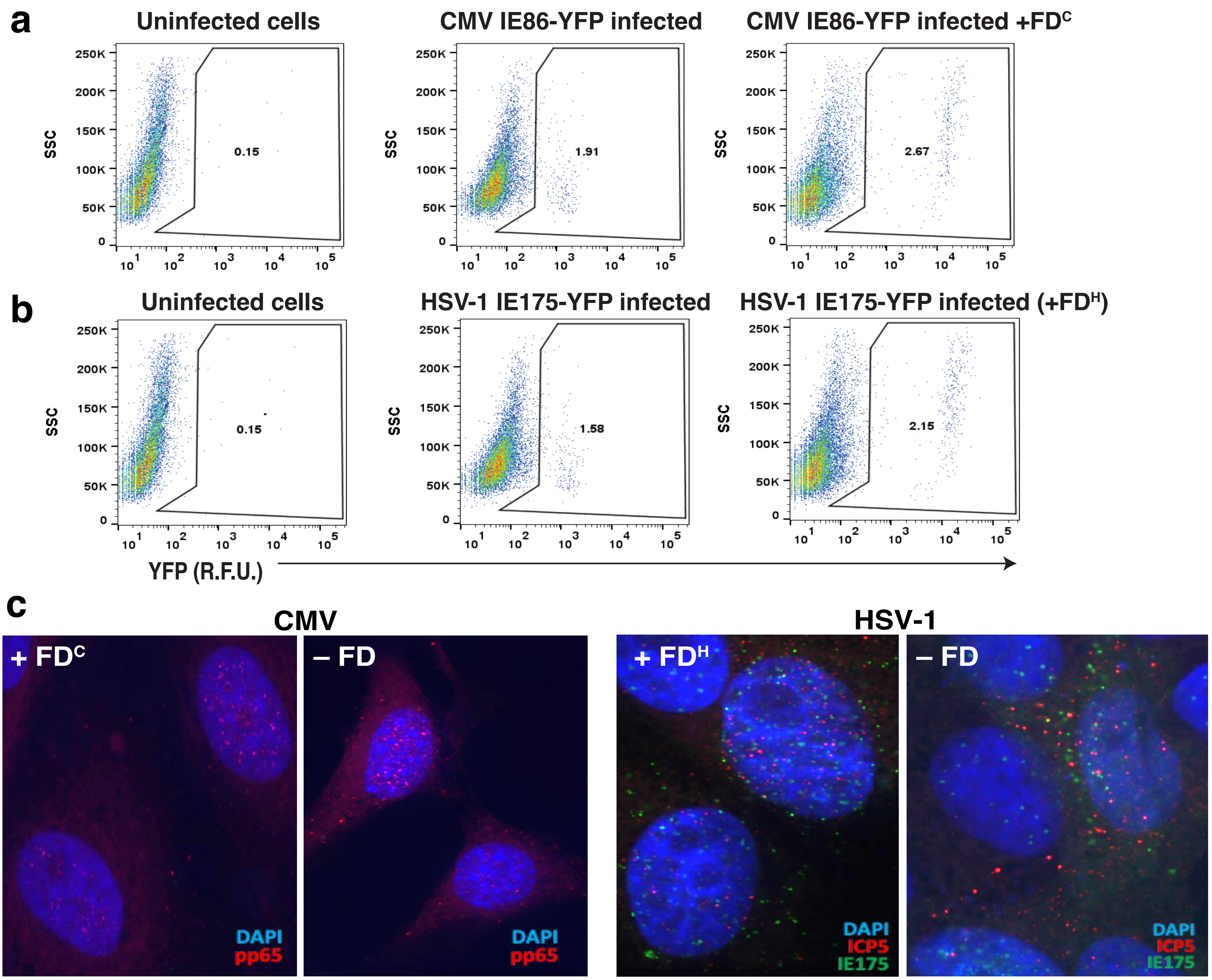

### Extended Data Figure 5 (Chaturvedi et al.)

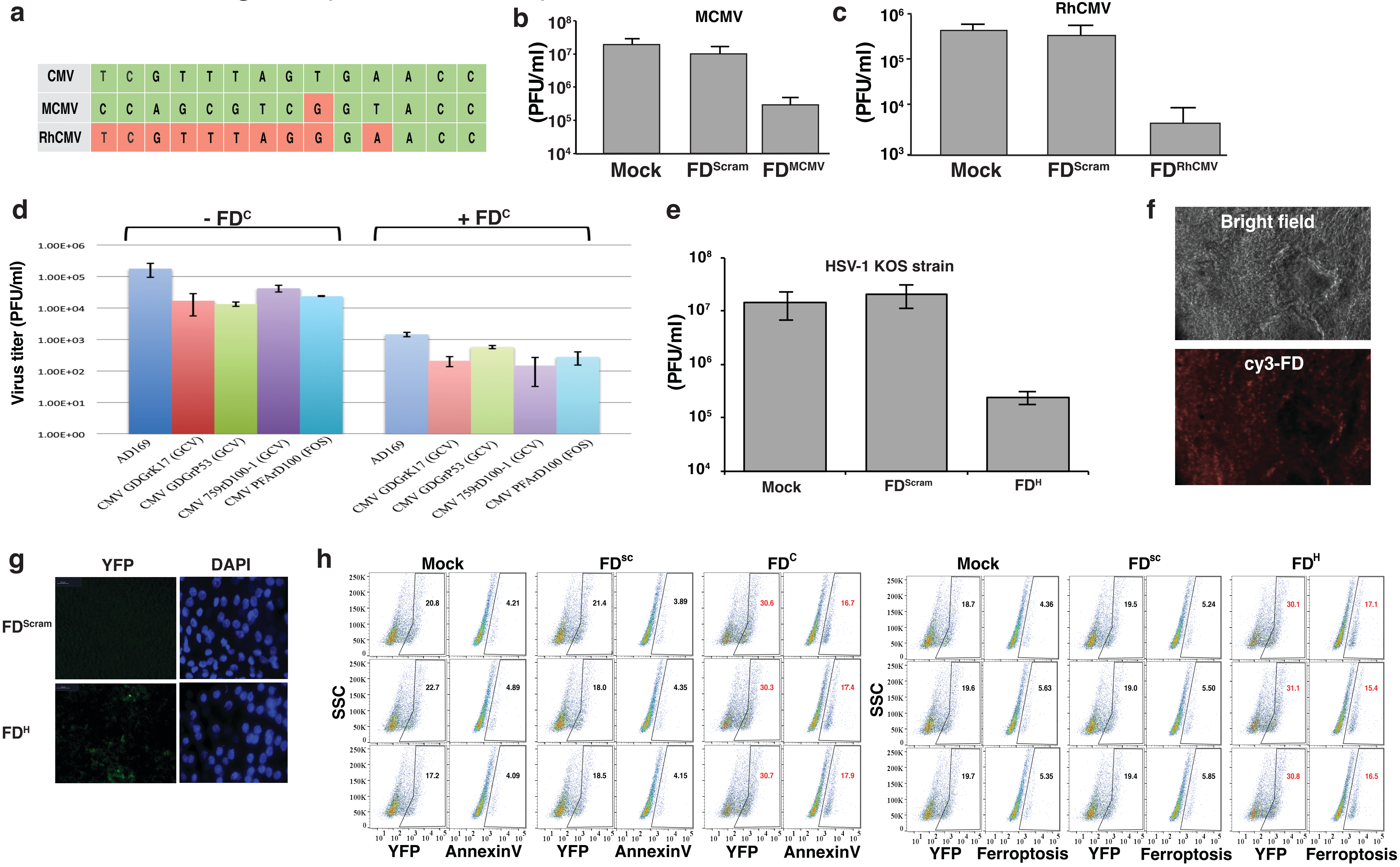

Extended Data Figure 6 (Chaturvedi et al.)

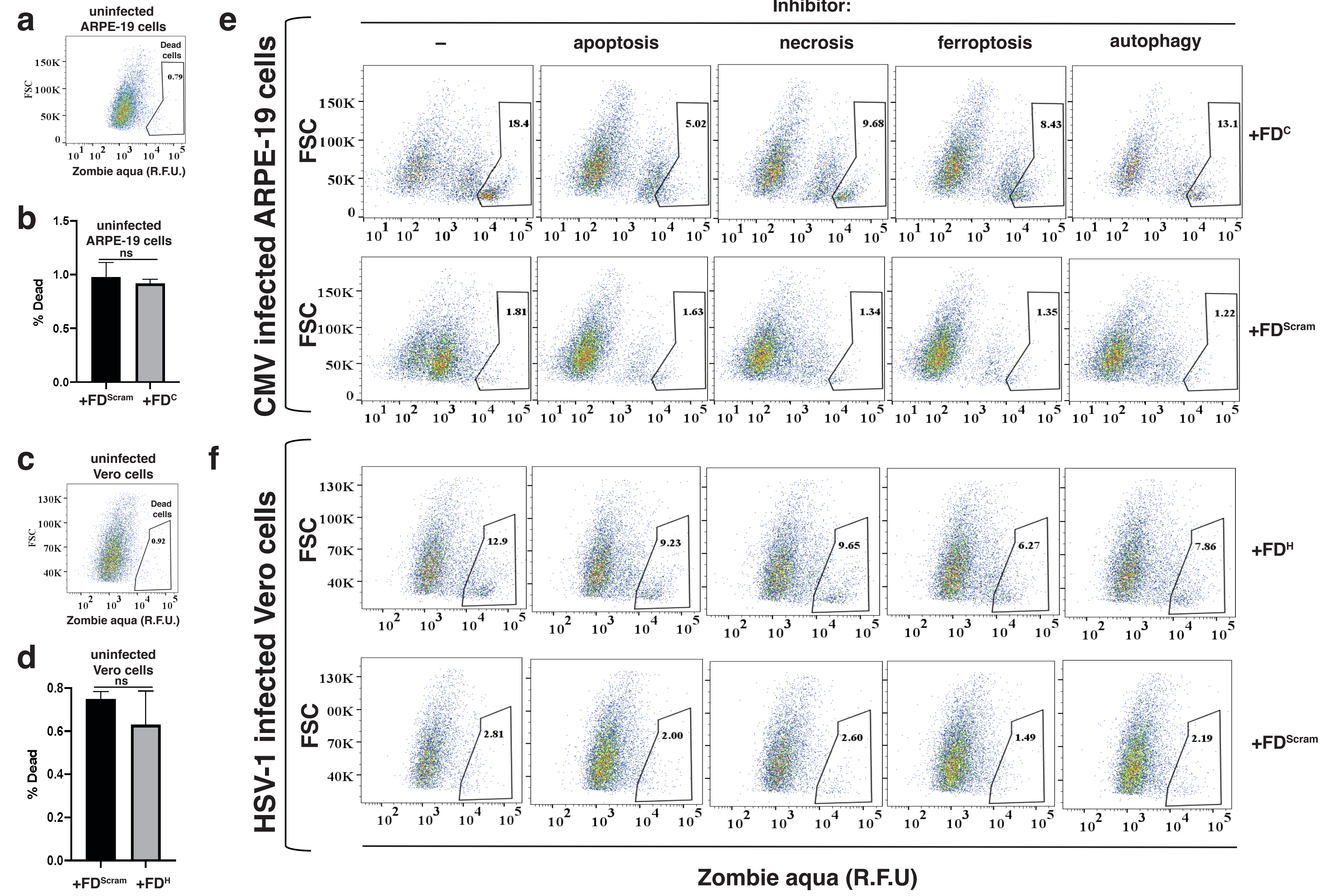

### Extended Data Figure 7 (Chaturvedi et al.)

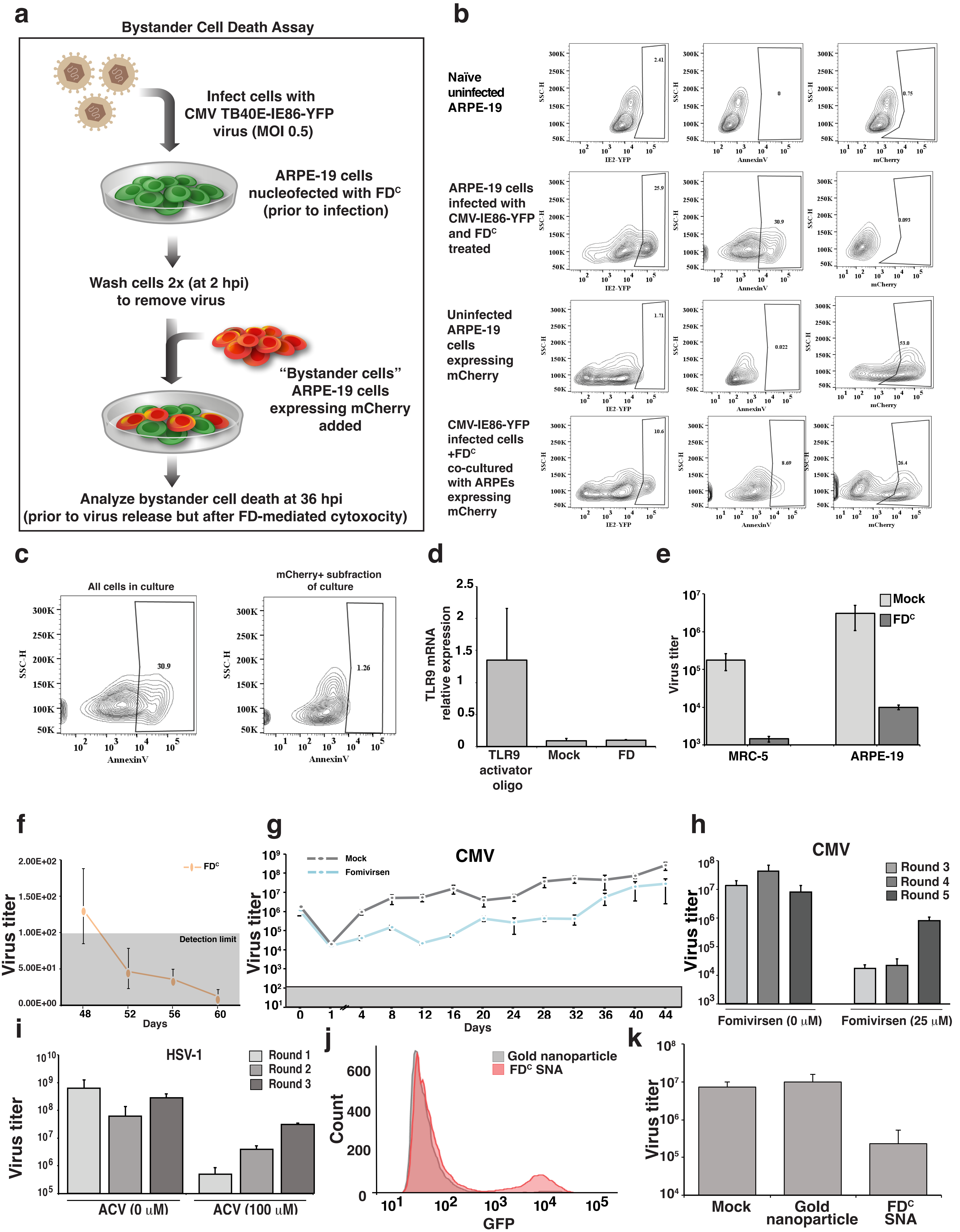

### Extended Data Figure 8 (Chaturvedi et al.)

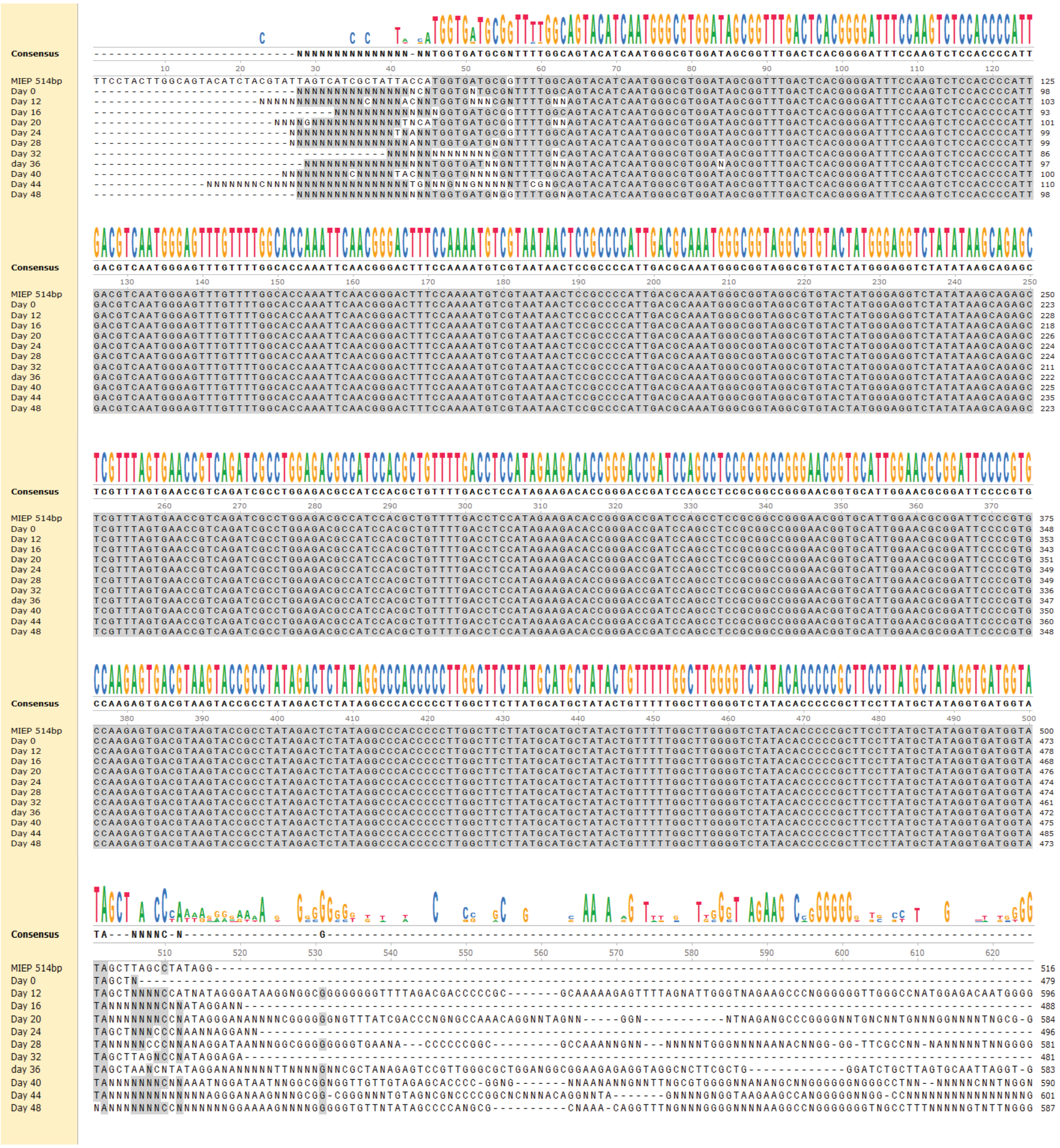

### Extended Data Figure 9 (Chaturvedi et al.)

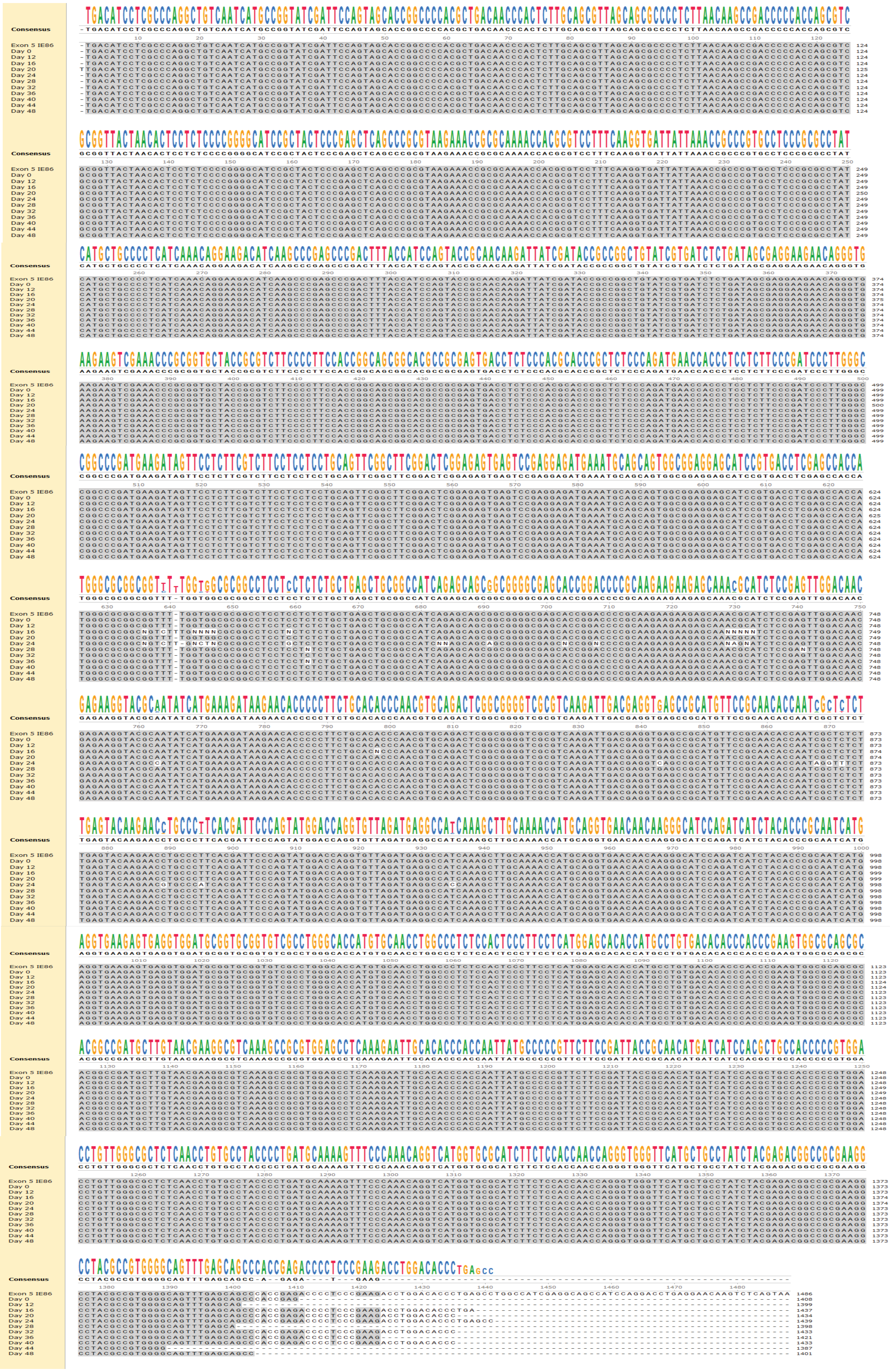

| No | Name | Primer Sequences (crs sequences highlighted in cyan; * denotes PS bonds) |
|---|---|---|
| 1 | crs for EMSA | Fw: GGTGGGAGGTCTATATAAGCAGAGCTCGTTTAGTGAACCGTCAGATCGCCTGGAGAC<br>Rw: GTCTCCAGGCGATCTGACGGTTCACTAAACGAGCTCTGCTTATATAGACCTCCCACC |
| 2 | Δcrs for EMSA | Fw: GGTGGGAGGTCTATATAAGCAGAGCCTAGGTAGTGAACCGTCAGATCGCCTGGAGC<br>Rw: GCTCCAGGCGATCTGACGGTTCACTACCTAGGCTCTGCTTATATAGACCTCCCACC |
| 3 | 14bp DNA | Fw: TCGTTTAGTGAACC<br>Rw: GGTTCACTAAACGA |
| 4 | 21bp DNA | Fw: AGATCGTTTAGTGAACCGTAC<br>Rw: GTACGGTTCACTAAACGATCT |
| 5 | 24bp DNA | Fw: CAGATCGTTTAGTGAACCGTACAC<br>Rw: GTGTACGGTTCACTAAACGATCTG |
| 6 | 26bp DNA | Fw: ACAGATCGTTTAGTGAACCGTACACG<br>Rw: CGTGTACGGTTCACTAAACGATCTGT |
| 7 | 28bp DNA | Fw: GACAGATCGTTTAGTGAACCGTACACGA<br>Rw: TCGTGTACGGTTCACTAAACGATCTGTC |
| 8 | 30bp DNA | Fw: AGACAGATCGTTTAGTGAACCGTACACGAT<br>Rw: ATCGTGTACGGTTCACTAAACGATCTGTCT |
| 9 | 34bp DNA | Fw: ATAGACAGATCGTTTAGTGAACCGTACACGATTA<br>Rw: TAATCGTGTACGGTTCACTAAACGATCTGTCTAT |
| 10 | 64bp DNA | Fw:TAATACGACTCACTATAGGGCGAATTGGAGCTCGTTTAGTGAACCGTCAGATCTCTAGAAGCTT<br>Rw:AAGCTTCTAGAGATCTGACGGTTCACTAAACGAGCTCCAATTCGCCCTATAGTGAGTCGTATTA |
| 11 | FD$^{C}$ | G*ACAGATCGTT*TAGTGAACC*GTACACG*A<br>T*CGTGTAC*GGTTCACTA*AACGATCTGT*C |
| 12 | FD$^{Scram}$ (for CMV) | Fw: A*TACCGCGTAA*CCAGAGGT*ATATAGGT*C<br>Rw: G*ACCTATAT*ACCTCTGG*TTACGCGGTA*T |
| 13 | FD$^{RhCMV}$ | Fw: G*ACAGATCGT*TTAGGGAAC*CGTACACG*A<br>Rw: T*CGTGTACG*GTTCCCTAA*ACGATCTGT*C |
| 14 | FD$^{MCMV}$ | Fw: G*ACAGACCA*GCGTCGG*TACCGTACACG*A<br>Rw: T*CGTGTACGGT*ACCGACGC*TGGTCTGT*C |
| 15 | Cy3-FD$^{C}$ | /5Cy3/G*ACAGATCGTT*TAGTGAACC*GTACACG*A<br>T*CGTGTAC*GGTTCACTA*AACGATCTGT*C |
| 16 | FD$^{C}$ (1x) | Fw: CGTTTAGTGAACCGCGCGCGATAGTACATAATCA<br>Rw: TGATTATGTACTATCGCGCGCGGTTCACTAAACG |
| 17 | FD$^{C}$ (2x) | Fw: CGTTTAGTGAACCGCGCGCGCGTTTAGTGAACCG<br>Rw: CGGTTCACTAAACGCGCGCGCGGTTCACTAAACG |
| 18 | FD$^{H}$ | Fw: CCG*AGGAC*GCCCCGATC*GTCCACACG*GAG<br>Rw: CTC*CGTGTGGAC*GATCGGGGC*GTCCT*CGG |
| 19 | FD$^{Scram}$ for HSV-1 | FW: GCA*GACCG*CTGGACGCG*CCAACACTC*CGG<br>RW: CCG*GAGTG*TTGGCGCGT*CCAGCGGTC*TGC |
| 20 | Cy3-FD$^{H}$ | Fw: /5Cy3/CCG*AGGAC*GCCCCGATC*GTCCACACG*GAG<br>Rw: CTC*CGTGTGGAC*GATCGGGGC*GTCCT*CGG |
| 21 | ODN 2216 FW | Fw: GGGGGACGATCGTCGGGGGG<br>Rw: CCCCCCGACGATCGTCCCCC |
| 22 | TLR9-qPCR | Fw: CCGTGACAATTACCTGGCCTTC<br>Rw: CAGGGCCTTCAGCTGGTTTC |
| 23 | MIEP 250bp crs | Fw: TTCCTACTTGGCAGTACATCTAC<br>Rw: CCTATAGGCTAAGCTATACCATC |
| 24 | IE86 Exon 5 | Fw: TGACATCCTCGCCCAGG<br>Rw: TTACTGAGACTTGTTCCTCAGGT |
| 25 | IE86-qPCR | Fw: TGACCGAGGATTGCAACGA<br>Rw: CGGCATGATTGACAGCCTG |
| 26 | IE175-qPCR | Fw: CCTATAGGCTAAGCTATACCATC<br>Rw: GTCTGACGGTCTGTCTCTGG |
| 27 | GAPDH-qPCR | Fw: TTCGACAGTCAGCCGCATCTT<br>Rw: CAGGCGCCCAATACGACCAAA |
